# *Streptococcus pyogenes* pharyngitis elicits diverse antibody responses to key vaccine antigens influenced by the imprint of past infections

**DOI:** 10.1101/2024.04.20.590388

**Authors:** Joshua Osowicki, Hannah R Frost, Kristy I Azzopardi, Alana L Whitcombe, Reuben McGregor, Lauren H. Carlton, Ciara Baker, Loraine Fabri, Manisha Pandey, Michael F Good, Jonathan R. Carapetis, Mark J Walker, Pierre R Smeesters, Paul V Licciardi, Nicole J Moreland, Danika L Hill, Andrew C Steer

## Abstract

Knowledge gaps regarding human immunity to *Streptococcus pyogenes* have impeded vaccine development. To address these gaps and evaluate vaccine candidates, we established a human challenge model of *S. pyogenes* pharyngitis. Here, we analysed antibody responses in serum and saliva against 19 antigens to identify characteristics distinguishing 19 participants who developed pharyngitis and 6 who did not. Pharyngitis elicited serum IgG responses to key vaccine antigens and a muted mucosal IgA response, whereas the 6 participants without pharyngitis had more pronounced IgA responses and minimal IgG responses. Serum IgG responses to pharyngitis in adult participants resembled those observed in children and were inversely correlated with the magnitude of pre-existing responses. While a straightforward correlate of protection was not evident, baseline antibody signatures distinguished clinical and immunological outcomes following experimental challenge. This highlights the influence of a complex humoral imprint from previous exposure, relevant for interpreting immunogenicity in forthcoming vaccine trials.

## Introduction

*Streptococcus pyogenes* is a highly adapted human-restricted pathogen with an immense global burden of communicable and non-communicable disease across a diverse clinical spectrum spanning acute infections and post-infectious syndromes with chronic disease outcomes^1^. *S. pyogenes* ranks in the global top 10 infection-related causes of death, with more than 500,000 deaths every year attributable to rheumatic heart disease^2^ and acute invasive infections^3^. In recognition of the clear unmet need, the World Health Organization Product Development for Vaccines Advisory Committee lists *S. pyogenes* as a global priority pathogen for new vaccine research and development^4^. While recent efforts have successfully reinvigorated the field^5,6^, the modern *S. pyogenes* vaccine development ecosystem remains fragile^7^.

Critically, there is no established human immune correlate of protection to predict efficacy of *S. pyogenes* vaccines^8^. The epidemiology of *S. pyogenes* infections strongly suggests that partial immune protection accumulates with repeated exposure through childhood^8–10^. However, unlike vaccine-preventable bacterial diseases caused by *Streptococcus pneumoniae*, *Haemophilus influenzae* and *Neisseria meningitidis*, there are no primary or acquired immunodeficiency syndromes classically associated with susceptibility to *S. pyogenes* infection from which the basis for naturally acquired protection may be inferred. The contribution of M protein serotype-or *emm* genotype-specific humoral immunity, initially established by streptococcal research pioneer Rebecca Lancefield^11^, remains uncertain^8^, with mixed findings from longitudinal cohort studies^12–15^, historical human challenge trials^16–19^, animal models^20^, and *in vitro* assays^21^.

In the early 20^th^ century, hundreds of thousands of children and adults were vaccinated for prevention of *S. pyogenes* clinical syndromes, all considered then as ‘scarlet fever’^22^. The last time efficacy of a *S. pyogenes* vaccine was evaluated in humans was in 1970s challenge trials in which parenteral and mucosal monovalent M protein vaccines were protective against homologous pharyngeal challenge, although a correlate of protection could not be demonstrated^17,18^. Since a contentious 1979 United States Food and Drug Administration ruling that slowed *S. pyogenes* vaccine research was revoked in 2006^23,24^, many M-protein and non-M protein antigens have proven to be protective in a variety of animal models^25^. Human immunogenicity for most of these antigens has been demonstrated in natural cohorts^12,15^, pooled human immunoglobulin products^26,27^, and a small number of early phase vaccine trials^28–32^. Still, by 2016, in the face of scientific, commercial, and regulatory barriers, *S. pyogenes* vaccines were considered ‘impeded vaccines’^33^.

As part of resurgent global *S. pyogenes* vaccine development efforts^5,6^, establishment of new human infection models has been prioritised as a platform for early efficacy evaluation to accelerate vaccine development^34^ and to explore immune responses. The CHIVAS-M75 trial established the world’s only modern *S. pyogenes* human infection model in healthy adult volunteers^35^. We have previously described a distinct systemic and mucosal cellular and cytokine signature of experimental human pharyngitis in CHIVAS-M75 participants^36^. Here, we aimed to interrogate longitudinal systemic, mucosal, functional, and binding antibodies to *S. pyogenes* in serum and saliva collected from 25 participants in the CHIVAS-M75 trial, before and after challenge, 19 of whom developed acute symptomatic pharyngitis and 6 who did not. These data lay the foundations for immune assessments in forthcoming trials to evaluate promising vaccine candidates^7^.

## Results

### Antibodies induced by challenge have modest activity in functional in vitro bacterial adhesion and opsonophagocytic assays

We first investigated mucosal and systemic functional responses against the *emm*75 *S. pyogenes* challenge strain^37^ using an established serum opsonophagocytic killing assay^38^ and an adapted bacterial adhesion assay^39^ (Fig. 1). Saliva collected before and 1-week after challenge affected bacterial adhesion to a pharyngeal cell line (Detroit 562), however there were no consistent changes related to challenge or between those who did and did not develop pharyngitis (Fig. 1B). We detected no serum opsonophagocytic activity against the challenge strain for 23 of 25 participants in samples from before and 1-month after challenge (Fig. 1C). Opsonophagocytic activity was observed at baseline in just two participants, one who subsequently developed pharyngitis (SN010) and one who did not (SN013). These were the only participants with increased opsonophagocytic killing after 1-month. Both had sustained pharyngeal colonisation with *S. pyogenes*, anti-streptolysin O seroconversion^35^, and increased inhibition by post-challenge saliva in the adhesion assay. In summary, neither functional assay correlated with clinical outcome.

**Figure 1.**
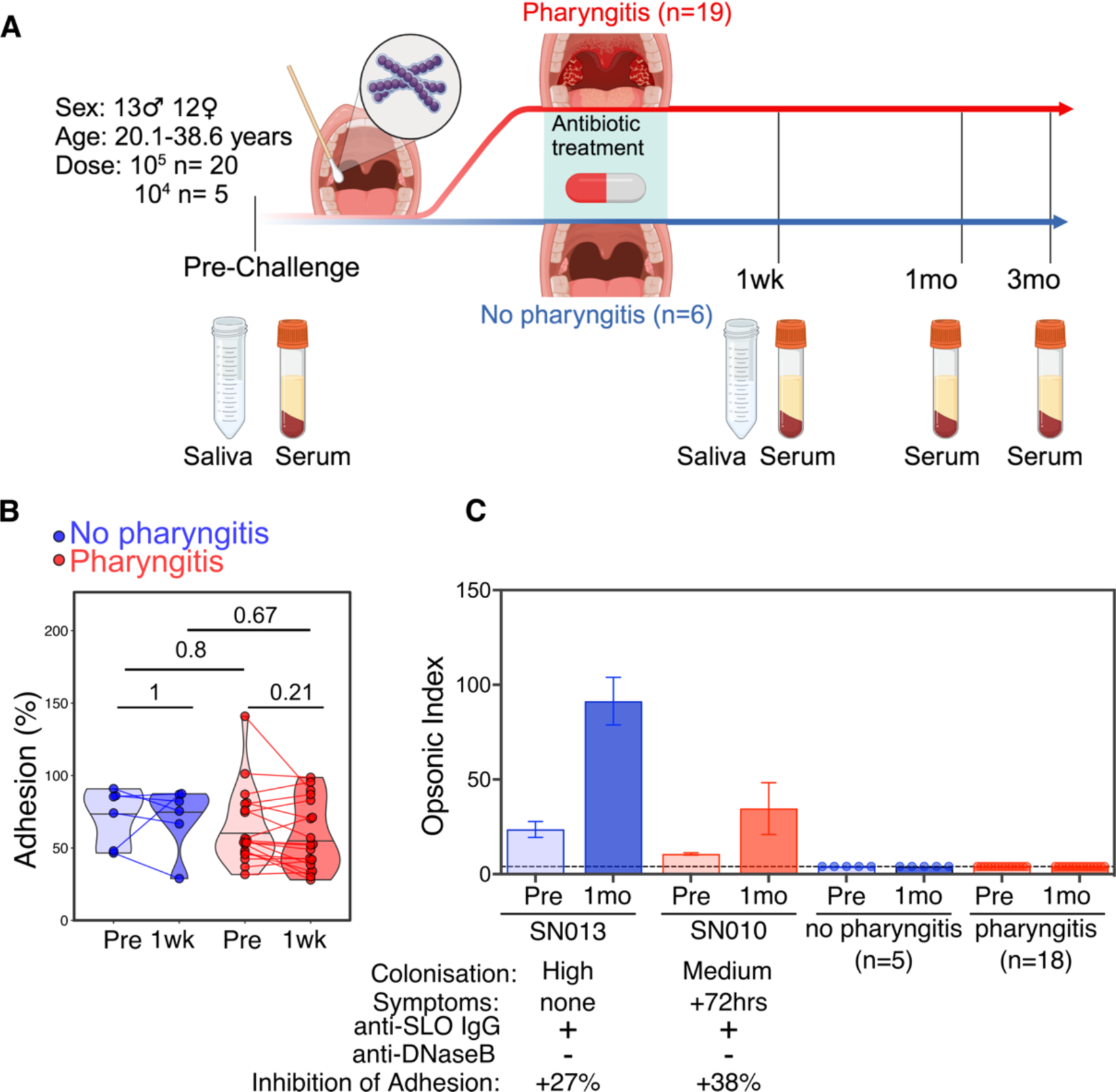
Experimental human *S. pyogenes* pharyngitis is not associated with new serum opsonophagocytic responses or inhibition of adherence by saliva. A) Schematic overview of challenge study timelines and samples analysed in this study. B) Bacterial adherence to Detroit 562 cells in the presence of saliva collected pre-challenge and 1 week after pharyngitis diagnosis (n=19) or discharge without pharyngitis (n=6). Bacterial burden determined by CFU of recovered bacteria and % adherence determined relative to bacteria-only controls. Responses for each group shown as median+range, with within group comparisons performed using paired Wilcoxon signed-rank tests and between group comparisons using Mann-Whitney tests, p-values are shown following false discovery rate (FDR)-adjustment for multiple comparisons. C) Serum opsonophagocytic killing assay results from pre-challenge and one-month post-challenge samples (n=25), highlighting results from 2 participants with a detectable opsonophagocytic response (SN013, SN010; mean and standard error of three technical replicates shown), and grouping the remainder of non-responders by clinical pharyngitis outcome (blue = non-pharyngitis, n=5; red = pharyngitis, n=18).

Opsonic index determined by the linear regression of the 2 dilutions closest to 50% killing using Opsititer software. Text descriptions below describe clinical and infection parameters for SN013 and SN010.

### Pharyngeal challenge induces heterogenous serological responses against key vaccine antigens

To evaluate humoral immune responses, we quantified IgG in serum and IgA in saliva against a panel of 17 antigens comprising vaccine candidates and known targets of *S. pyogenes* immunity (Fig. 2A, Table S1). IgG and IgA responses were highly variable across antigens for all participants at pre and post-challenge time-points (Fig 2B), including inconsistent responses against protein antigens and related derivative peptides, such as the M75 protein and its derivative antigens: the N-terminal M75-HVR peptide and C-repeat peptides p145, J8, and P*17 (Figs S2, S3). When analysed as fold-change relative to pre-challenge titres, participants who developed pharyngitis had increased post-challenge serum IgG responses against several key vaccine antigens (SpyCEP, SLO, ScpA, GAC) and a trend towards lower post-challenge saliva IgA responses, especially for GAC and M75-HVR (Figs 2C, 2F). Conversely, among the smaller group of 6 participants who did not develop pharyngitis, there were trends towards lower serum IgG and higher saliva IgA post-challenge responses. These trends were antigen-specific, as total IgA concentration in saliva was not affected by challenge or pharyngitis (Fig S1).

**Figure 2:**
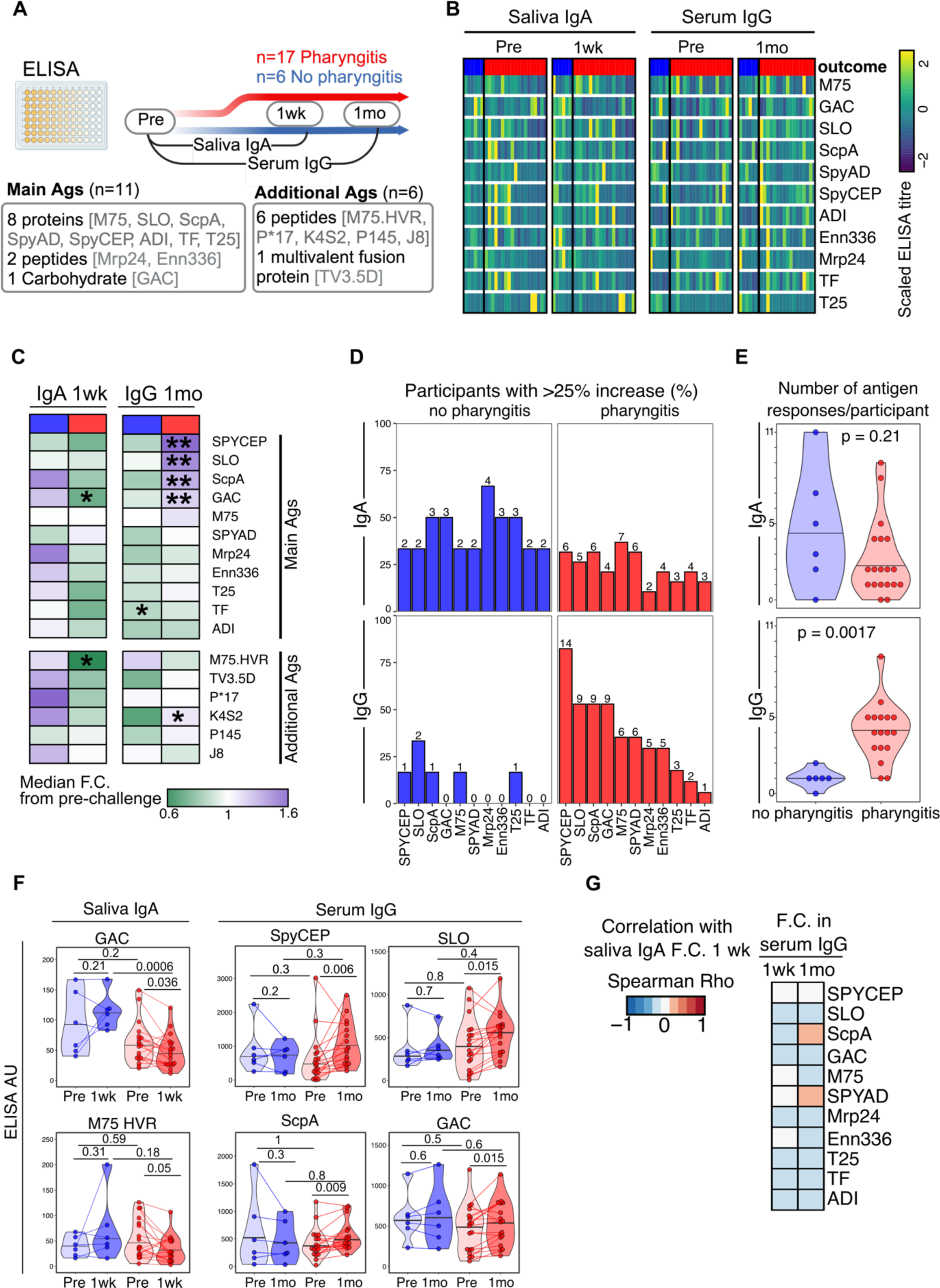
Experimental pharyngeal challenge with *Streptococcus pyogenes* induces mucosal and systemic antibody responses against major vaccine antigens. A) Schematic overview of the time-points and antigens studied to detect IgG from serum and IgA saliva. ‘Main’ Ags include leading vaccine candidates and key virulence factors, and ‘Additional’ Ags include peptides or domains of proteins included among Main Ags (M protein, SpyCEP, T antigen).B) Heatmap showing relative abundance of IgA and IgG responses pre and post-challenge at indicated time-points in pharyngitis and non-pharyngitis participants for 11 main antigens. ELISA titres are shown with responses scaled across IgA and IgG separately. Pharyngitis outcome indicated as coloured bar (red = pharyngitis, blue = no pharyngitis).C) Heatmap showing the median fold-change observed relative to baseline at 1 week for IgA, and 1-month for IgG with associated p-values from paired Wilcoxon-signed rank test after FDR-adjustment for 11 main antigens and 6 additional peptide or vaccine antigens. D) The percentage of pharyngitis and non-pharyngitis participants that showed a 25% increase in antibodies post-challenge, with 1 week for IgA and 1-month for IgG. Numbers indicate how many participants responded to each antigen. E) A tally of the number of antigens to which participants showed a 25% increase in IgA and IgG (shown in D), with participants split by pharyngitis outcome. P-value determined from Mann-Whitney U-test. F) Representative examples of antigens that showed significant change after challenge (as shown in C) expressed in ELISA arbitrary units (AU). G) Spearman correlation of fold-change in saliva IgA at 1 week with fold-change in serum IgG at 1 week and 1-month post-challenge. All comparisons were p>0.1. Within group comparisons performed using paired Wilcoxon signed-rank test, and groups compared using Mann-Whitney test, all FDR-adjusted for multiple comparisons.

We sought to determine the frequency of responses to each antigen and if this differed by pharyngitis outcome. A ‘response’ was defined as an increase of 25% or greater above pre-challenge titres, a threshold previously applied for IgG responses to other bacterial pathogens^41,42^, and used here to capture the breadth of antigens recognised by the modest antibody response induced by a single infection. Serum IgG responses were more common among participants with pharyngitis, whereas antigen-specific saliva IgA responses occurred with similar or higher frequency among participants without pharyngitis (Fig. 2D). Consistent with the fold-change findings, a response was observed most frequently against 4 vaccine candidate antigens (SpyCEP, SLO, ScpA and GAC). The breadth of the serum IgG response was higher among participants with pharyngitis, with a median response to 4 antigens, compared to one antigen among participants without pharyngitis (Fig 2E). When analysing saliva, participants with pharyngitis responded to a median of 2 antigens, while there was a trend towards a greater breadth of IgA responses among participants without pharyngitis, including one participant with a saliva IgA response to 11 different antigens. Overall, the most robust post-challenge differences among participants after pharyngitis were decreased saliva IgA for GAC and M75-HVR and increased serum IgG for SpyCEP, SLO, ScpA, and GAC (Fig 2F, all other antigens Figs S2-S4). The opposing trends observed for IgG and IgA responses led us to consider if these responses were somehow related. However, there was no correlation observed between fold-change of saliva IgA responses and serum IgG responses at either 1 week or 1-month (Fig 2G).

### Experimental S. pyogenes pharyngitis induces a long-lived humoral immune response

For the 4 antigens associated with the most frequent serum IgG responses (SpyCEP, SLO, ScpA, GAC), we proceeded to use a multiplex bead-based assay to determine the absolute concentration of antigen-specific serum IgG^43,44^. Responses to SpnA and DnaseB were also analysed because they can be used clinically (with SLO) as evidence of recent *S. pyogenes* infection^45^. For all 6 antigens, participants with a 25% increase in IgG concentration (the ‘responders’) all developed pharyngitis, except for a single participant who did not develop pharyngitis and was a responder to ScpA (Fig 3B). The increase in IgG to all antigens in responders occurred as early as the 1-week outpatient visit and was maintained out to 3 months indicating that when antibodies were induced, they were durable for several months. At each time-point, the absolute IgG concentration for responders was equivalent to non-responders, except for IgG against SpnA which had a higher median concentration for non-responders at all time-points. There was a trend towards higher pre-challenge titres in non-responders, most evident for SpyCEP, SpnA and SLO.

**Figure 3:**
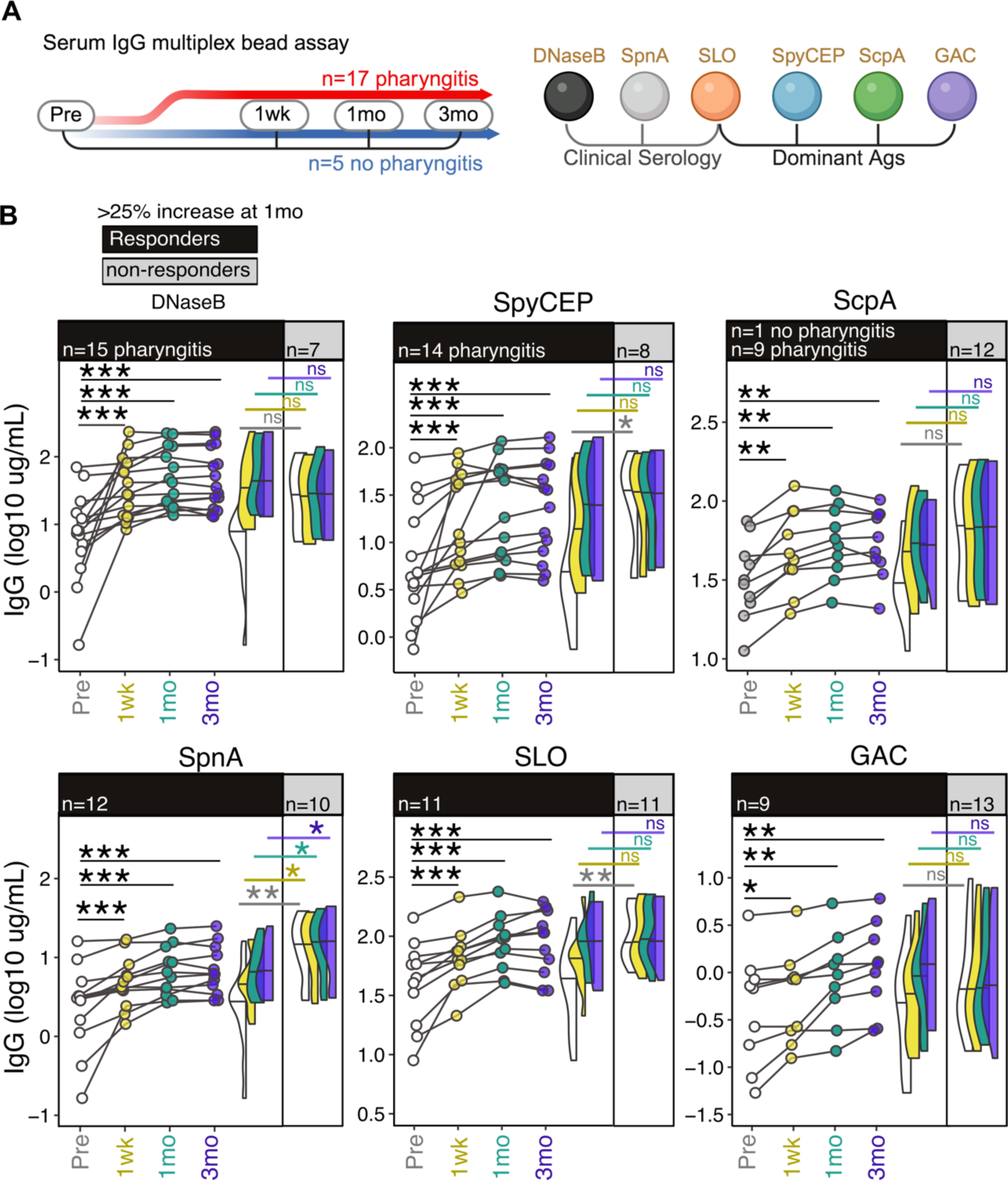
Antibody responses induced by experimental pharyngeal challenge with *Streptococcus pyogenes* are maintained for at least 3 months. A) Schematic overview of time points and antigens measured by a multiplex bead-based assay for IgG antibodies in sera. B) IgG responses for 6 antigens at indicated time points with the cohort split into those who showed a 25% increase in IgG at 1-month post-challenge (‘responders’) and those that did not (‘non-responders’). For ‘Responders’, each dot represents an individual and lines connect paired samples. The violin plots summarise the distribution (median+range) of responses across time-points for each group, with colour corresponding to time-point. Analysis included 22 participants with samples available from all 4 time-points. P-values were determined for within-group paired sample comparisons using Friedman’s test followed by Wilcoxon signed-rank test with FDR-adjustments, and for between group comparisons using Kruskal-Wallis and Dunn’s multiple comparison testing.

### Antibody responses to experimental human S. pyogenes pharyngitis in adults resembles responses to natural infection in children

Current *S. pyogenes* initiatives are focussed on advancing a paediatric product. One concern regarding human challenge research is that immune responses in healthy adult participants may not be generalisable to children^34^. We compared serum IgG responses from the multiplex bead-based assay for the CHIVAS-M75 adult participants with previously published data for healthy children (n=15) and children one month following microbiologically-confirmed *S. pyogenes* pharyngitis (n=22) from New Zealand (aged 5 to 14 years)^46^. IgG concentrations were equivalent among pre-challenge adult samples and healthy children for all antigens except ScpA (Fig 4). Post-challenge adult samples had equivalent IgG concentrations for all 6 antigens to convalescent serum from children, with a trend towards elevated responses in the paediatric samples for SpnA only. These results provide some reassurance that pre-existing serum antibody concentrations and the magnitude of post-challenge responses in adult human challenge participants resemble those observed before and after pharyngitis in children.

**Figure 4:**
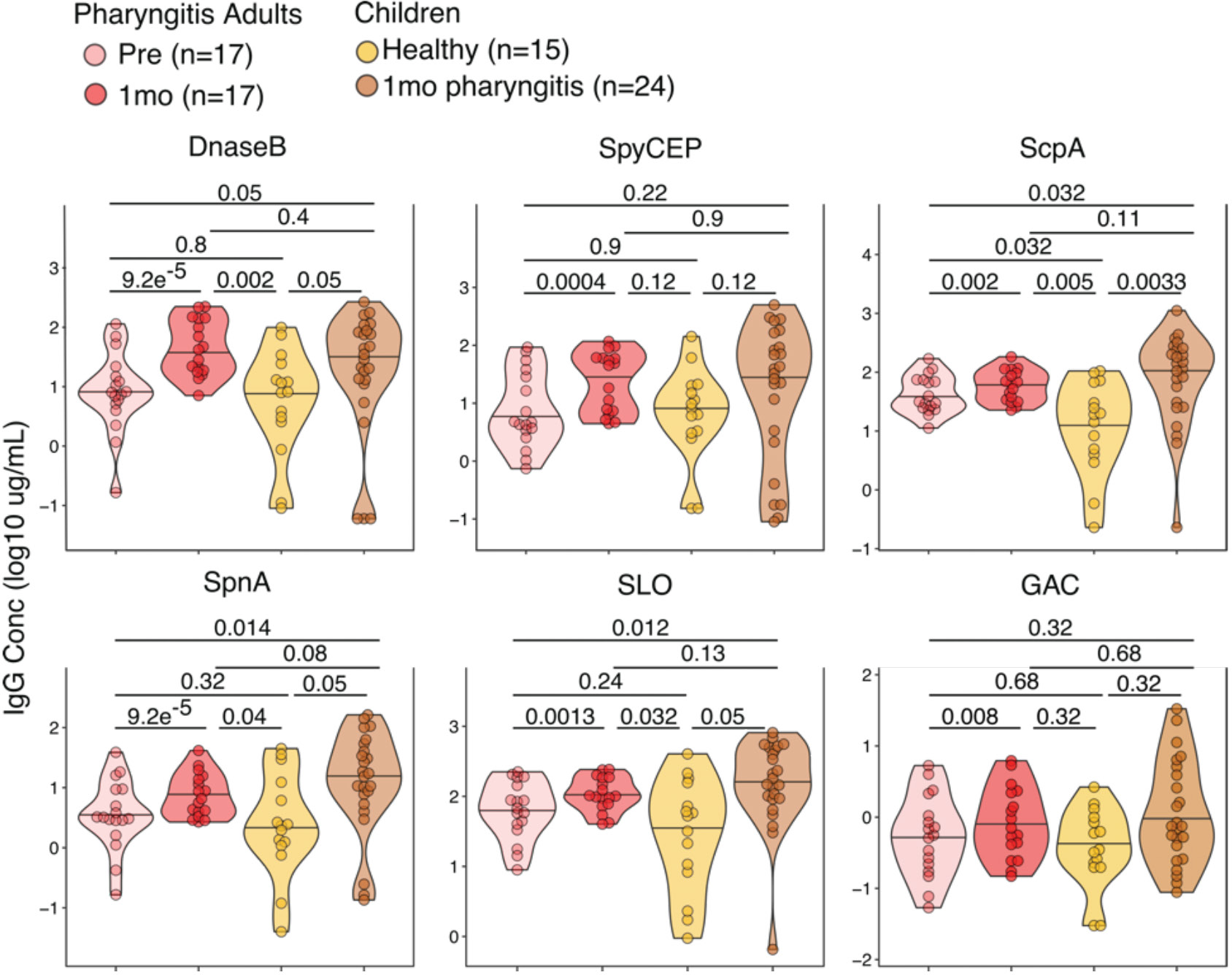
Antibody responses to experimental human *Streptococcus pyogenes* pharyngitis in healthy adults are similar to natural responses in children. Serum IgG concentrations measured by multiplex bead-based assay to 6 antigens among challenge participants pre and 1-month post-challenge was compared to healthy children and children > 1-month after acute pharyngitis (aged between 5 and 14 years)^46^. P-values determined using Dunn’s multiple-comparison testing, or Wilcoxon signed-rank test for paired data.

### Pre-existing humoral immunity inversely correlates with the post-challenge response

We sought to explore the higher pre-challenge IgG responses in ‘non-responders’. For 5 out of 6 antigens measured with the multiplex bead-based assay, there was a significant inverse correlation between pre-challenge IgG concentration and antibody fold-change by the 1-month outpatient visit, with a similar trend observed for GAC. Participants who did not develop pharyngitis were enriched among samples with higher pre-challenge serum IgG levels and minimal or negative post-challenge fold-change (Fig 5A). This inverse correlation was consistent for the 4 antigens measured by both multiplex bead-based assay and ELISA. For anti-M75, measured by ELISA, there was also a significant and similar inverse correlation (Fig 5B), although participants did not cluster distinctly based on outcome. Pre-challenge saliva IgA was inversely correlated with the fold-change in IgA by the 1-week outpatient visit for ScpA. In contrast to the IgG finding, participants without pharyngitis were enriched among samples with lower pre-challenge IgA and positive post-challenge fold-change (Fig 5C). Correlation coefficients for pre-versus post challenge IgG and IgA responses, measured by ELISA and multiplex-bead based assay, were negative for the majority of antigens tested. These data suggest that pre-existing antibodies may affect mucosal and systemic antibody responses to *S. pyogenes* infection, either by direct interference or as a surrogate for immune memory.

**Figure 5:**
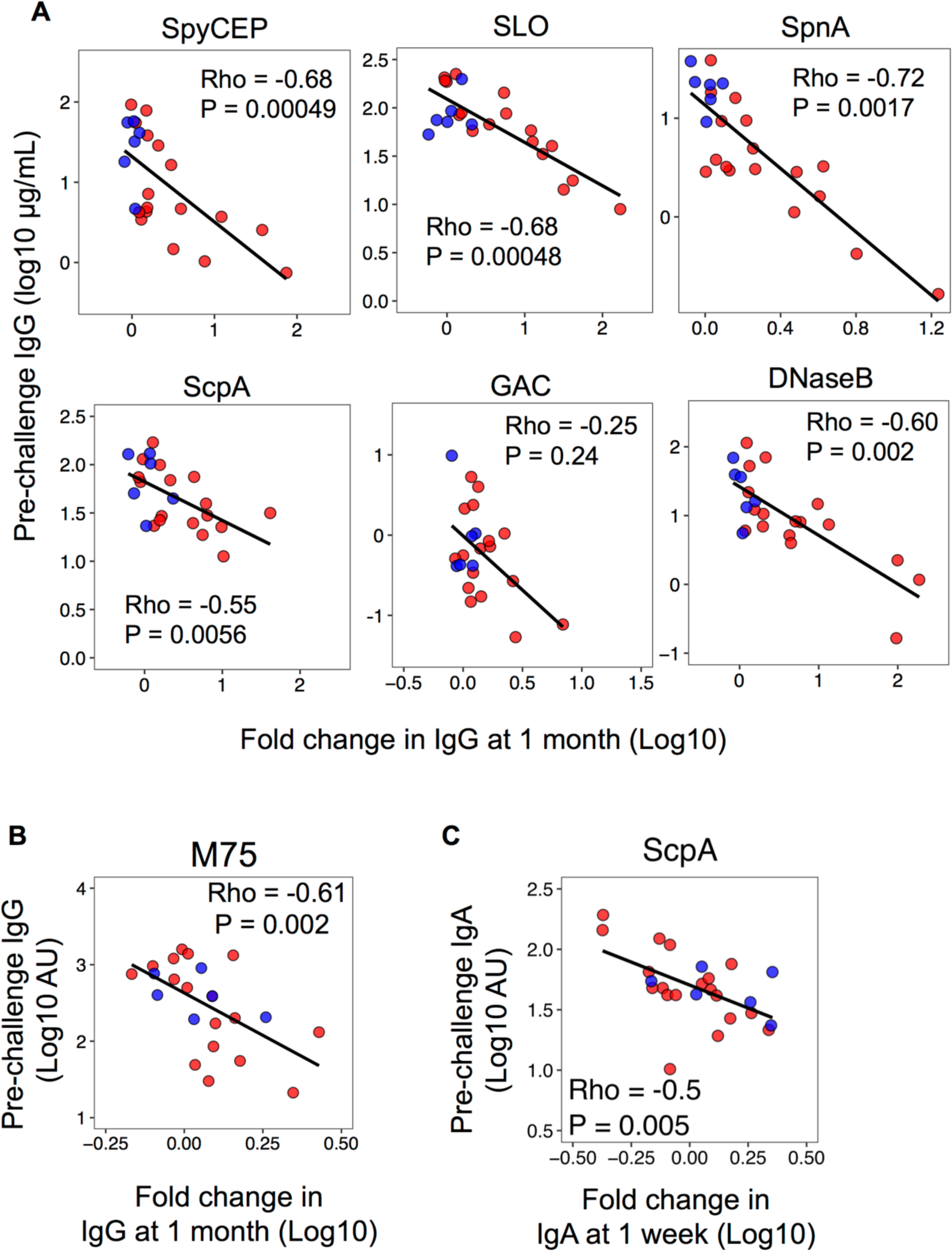
Pre-challenge baseline titres negatively correlate with post-challenge antibody responses. A) Correlation between IgG responses to 6 antigens measured by multiplex bead-based assay in samples from pre-challenge with those at 1-month post-infection (pharyngitis n=17, no pharyngitis n=6). B) Correlation between anti-M75 IgG responses measured by ELISA from pre-challenge with those at 1-month post-challenge (pharyngitis n=17, no pharyngitis n=6). C) Correlation between saliva anti-GAC IgA measured by ELISA in samples from pre-challenge and 1 week post-challenge (pharyngitis n=19, no pharyngitis n=6). Coefficient and p-value determined using Spearman’s method with linear regression line included to indicate linear trends. Each dot represents an individual participant with the pharyngitis group shown in red and non-pharyngitis in blue.

### Unbiased clustering reveals distinct serological signatures associated with experimental human pharyngitis and the humoral response to challenge

To investigate whether clinical outcome was associated with distinct serological features we used non-metric multidimensional scaling incorporating serum IgG data for 19 antigens (Table S1) from pre-challenge and the 1-month outpatient visit to generate an unbiased view of each participant’s serological signature. We analysed the IgG data to assess the correlation between each individual’s responses to all other individuals across the 19 antigens. Then, we used multidimensional scaling to simplify these complex relationships into two dimensions. Participants clustered according to pharyngitis outcome, suggestive of a pre-challenge serological state associated with protection from pharyngitis (left panel Fig 6A and 6B). For most participants, pre and post-challenge points were clustered closely together (middle panel Fig 6A), consistent with previous *S. pyogenes* exposures eliciting a distinct individual serological imprint. Where there was a change in multidimensional space coordinates for participants in response to challenge, this movement was predominantly left-to-right along Dimension 1 (middle panel Fig 6A) and was almost exclusively limited to the pharyngitis group (right panel Fig 6B). This suggests that, after challenge, some participants acquired a serological signature closer to the pre-challenge state of participants who did not develop pharyngitis. Dimension 1 was correlated with high serum IgG responses against GAC, SpyCEP, ScpA, K4S2, DnaseB, and SpnA, and low IgG responses to several antigens including J8, ADI and TF (Fig 6C). In keeping with this, participants with a left-to-right post-challenge shift along Dimension 1 were predominantly those we previously identified as IgG ‘responders’ against 2 or more of SpyCEP, GAC, SLO, and ScpA (Fig. 6A middle and right), indicating that responses to these antigens were major drivers of the clustering. Together, these data suggest that while individual human *S. pyogenes* serological signatures are highly variable, there are common features that may be associated with protection against pharyngitis.

**Figure 6.**
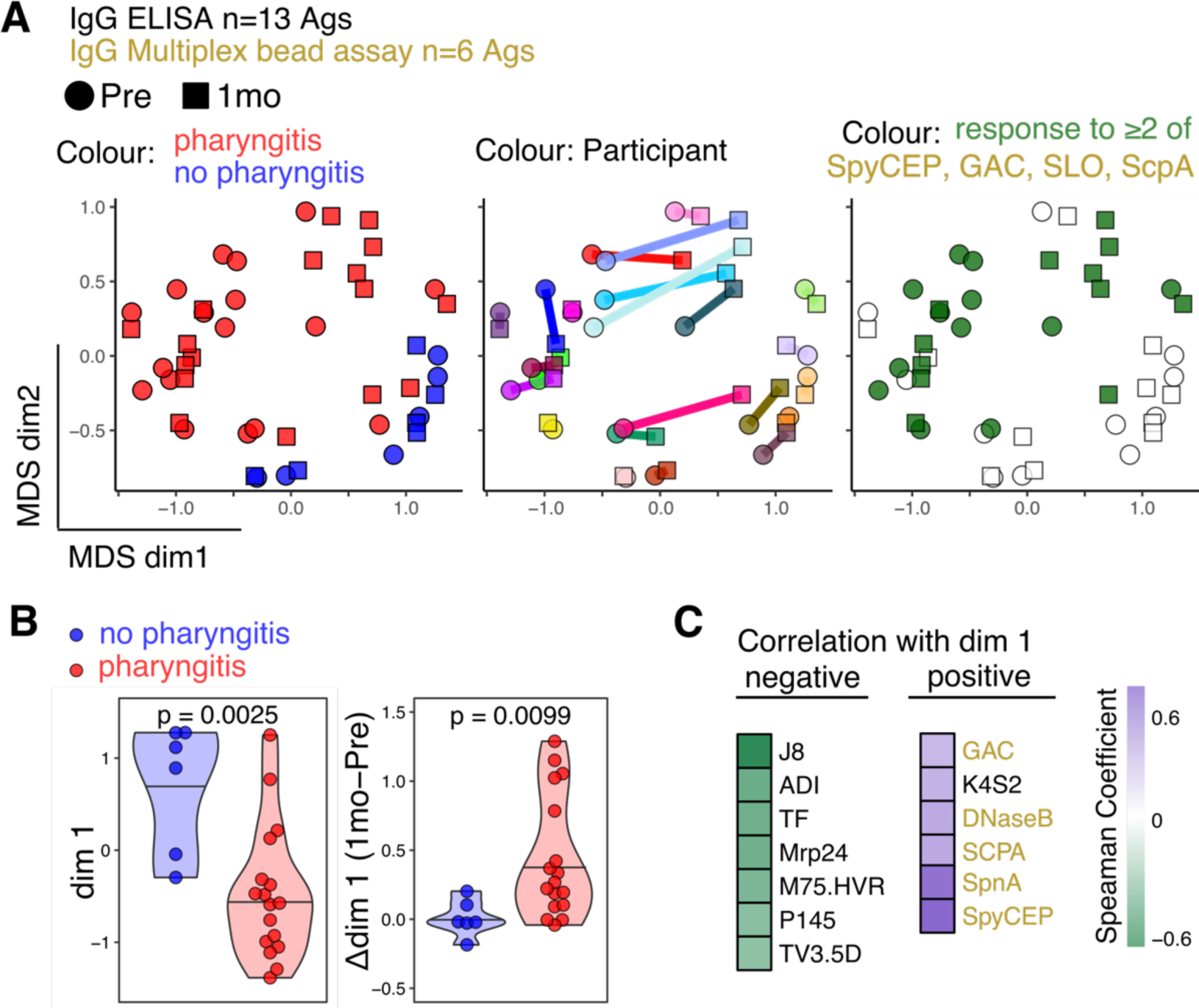
Serological signatures differ by clinical outcome following experimental pharyngeal human challenge with *S. pyogenes* and are shaped by infection. A) non-metric multidimensional scaling (MDS) was performed using IgG responses to 19 antigens for 23 individuals. Each point represents a sample, with shape denoting time point and coloured to indicate pharyngitis outcome (left), paired samples (centre), or responses to 2 or more of the dominant antigens SpyCEP, SLO, ScpA, or GAC. B) Violin plots of first MDS dimension values pre-challenge (left), or at 1-month with pre-challenge values subtracted (right), between pharyngitis and non-pharyngitis groups. P-value determined by Mann-Whitney U-test. C) Spearman correlation analysis between MDS dimension 1 and IgG responses, only antigens with significant correlation shown (p<0.01 after FDR-adjustment). Antigens with results from ELISA assays shown in black font and from multiplex bead-based assays shown in gold font.

### Baseline antibody levels are associated with clinical outcome and severity of experimental human S. pyogenes pharyngitis

We correlated 37 baseline antibody variables to 8 clinical features to investigate how pre-existing humoral immunity related to pharyngitis outcome and severity (symptoms, signs, time to diagnosis, intensity of bacterial colonisation) (Fig 7A)^35^. Ten variables significantly correlated with one or more clinical features (Fig 7B). While there were variable correlations between different antigens and clinical variables, for most antigens the directionality of correlation was strongly concordant. Broadly, baseline serum IgG against 3 antigens often described as ‘cryptic’ (ADI, TF^47^, J8^48^) correlated with severity of infections, whereas baseline antibodies against 7 other antigens were inversely correlated with severity: serum IgG against K4S2, DnaseB, SPyAD, SpyCEP, SpnA, T25, and saliva IgA against p145.

**Figure 7:**
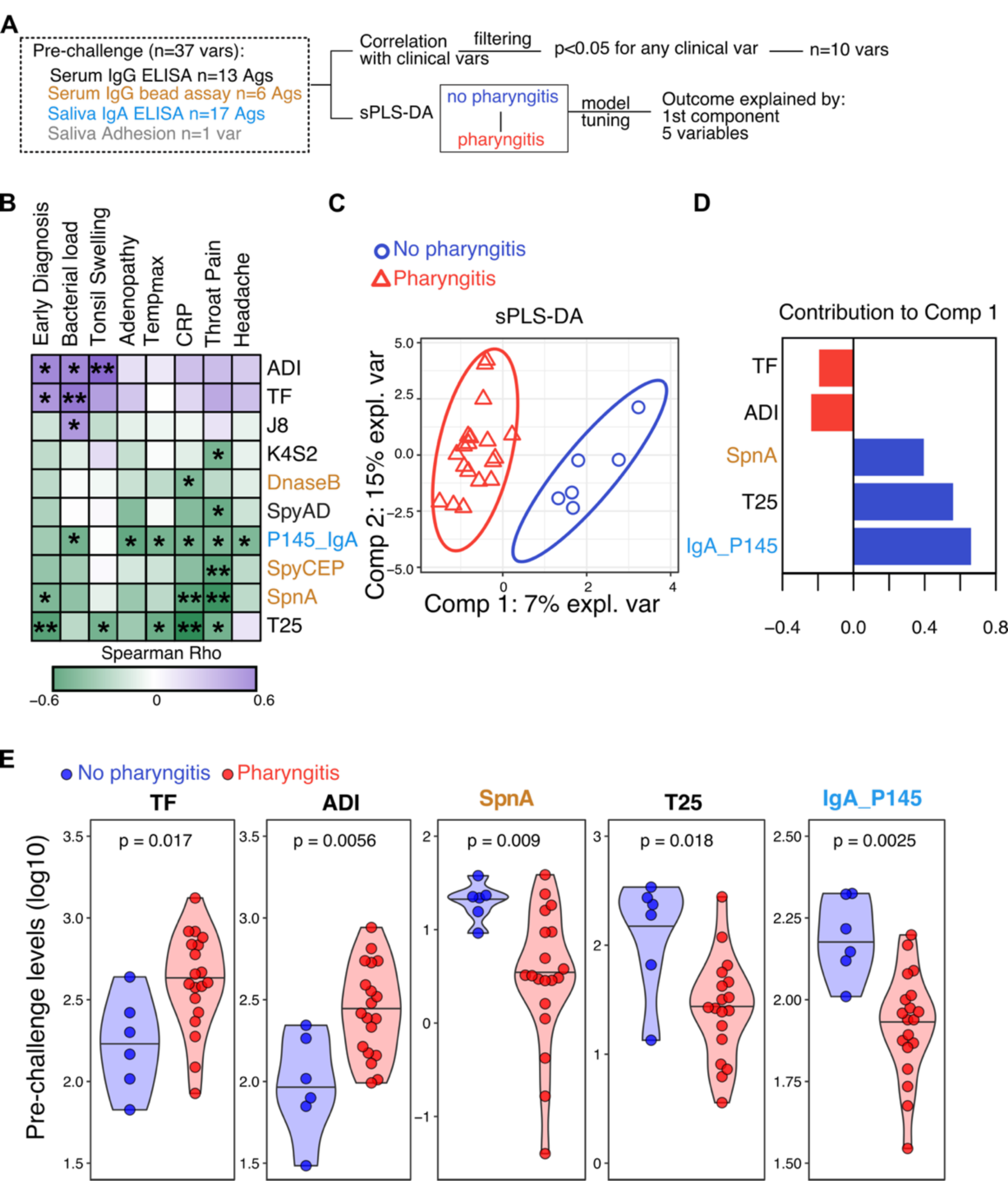
Baseline antibody titres are associated with clinical pharyngitis outcome following experimental human challenge with *Streptococcus pyogenes*. A) Schematic overview of methods and variables included in correlation analysis and sparse partial least squares discriminant analysis (sPLS-DA). B) Heatmap of correlation coefficients between various clinical variables and pre-challenge antibody levels. Antigens shown are those that showed significant correlation for one or more clinical variables. Asterisks denote p-values *<0.05, **<0.01. C) Samples distributed across 2 dimensions from sPLS-DA. D) Contributions of top 5 variables to sPLS-DA dimension 1, with variables in red higher in the pharyngitis group and variables in blue higher in the non-pharyngitis group. E) Pre-challenge antibody responses against the top 5 variables from sPLS-DA, differentiating between participants who subsequently developed pharyngitis (red) and those who did not (blue). P-value determined using Mann-Whitney Test. Antigens with results from ELISA assays shown in black (IgG) and blue (IgA) text and from multiplex bead-based assays (IgG) shown in gold text.

We then used sparse Partial Least Squares Discriminant Analysis (sPLS-DA) to discern which among the 37 immune variables most effectively distinguished between participants who did and did not develop pharyngitis. Analysing these variables in a multidimensional space, the sPLS-DA model pinpointed 5 variables that most significantly explained the immunological distinctions related to pharyngitis outcomes. Furthermore, the model revealed a single dimension along which participants were distinctly clustered according to their pharyngitis status (Fig 7C). The 5 pre-challenge variables that contributed to this first dimension and were associated with not developing pharyngitis were low serum IgG to TF and ADI, high serum IgG to SpnA and T25, and high saliva IgA to p145 (Fig. 7D), which were all robustly different between pharyngitis and non-pharyngitis groups (Fig. 7E). Together, these data suggest that the clinical outcome following *S. pyogenes* challenge is influenced by pre-existing humoral immunity and that systemic responses to some antigens may discriminate individuals with increased susceptibility to symptomatic pharyngitis.

## Discussion

This study of mucosal and systemic antibody responses in the CHIVAS-M75 human challenge trial advances our understanding of immunity against *S. pyogenes* and underlines the potential for the human model to accelerate vaccine development. Pharyngeal challenge induced heterogenous and highly individualised patterns of saliva IgA and serum IgG responses, with durable responses against conserved vaccine antigens (e.g. SpyCEP, ScpA, SLO, GAC). Serum IgG responses in healthy adult challenge participants were comparable to responses in children following natural pharyngitis. While a straightforward correlate of protection was not evident, baseline antibody signatures, reflecting the imprint of past infections, were identified that distinguished clinical outcomes following experimental challenge. Pre-existing antibodies inversely correlated with the magnitude of IgG induced by challenge, exemplifying the complex interactions between a highly adapted ubiquitous pathogen and its only natural host, with important ramifications for understanding vaccine immunogenicity.

Our study highlights the limitations of currently available functional serology assays, as the serum opsonophagocytic killing and salivary bacterial adhesion assays provided limited insight into the basis of susceptibility and protection. It is possible that the pre-screening and exclusion of individuals with high serum *emm*75-specific IgG from the CHIVAS-M75 trial^19,35^ may have contributed to the lack of opsonophagocytic killing observed. However, the M75 protein of the *emm*75 challenge strain was not the target of opsonisation by pooled human immunoglobulin (in the same assay) in a recent study^21^. These findings are consistent with 1970s human studies in which induction of a type-specific serum bactericidal response by M protein vaccines was not a reliable correlate of protection, which was observed against symptomatic pharyngitis (parenteral and mucosal vaccines) and colonisation (mucosal vaccine only)^16–18^. Expanding the scope of the modern human challenge model to include other *emm*-type strains will enable further evaluation of the opsonophagocytic killing assay as a correlate of protection. For multi-component vaccines including secreted virulence factors (e.g. SpyCEP, SLO), additional assays may be required to capture the diversity of functional antibody responses^49,50^.

Acute symptomatic *S. pyogenes* pharyngitis is characterised by a combined humoral and cellular immune response to contain and clear the focal infection^36^. The human model enabled simultaneous sequential assessment of IgA in saliva and IgG in blood. Antibody responses in these compartments did not correlate in magnitude or antigen-specificity, strongly suggesting they are produced and regulated independently. Overall, individual antibody responses were highly variable, as in previous longitudinal cohort studies of children^12,15,51^. Whilst every tested antigen was immunogenic in at least one participant, serum IgG responses were predominantly raised against conserved antigens with a role in evading human immunity (e.g. SpyCEP, SLO, ScpA), while responses to the M protein, conventionally considered as immunodominant, were modest and infrequent. In saliva, IgA responses were most pronounced against surface-bound antigens contributing to adhesion and invasion (e.g. M and M-like-proteins) and were more common among the 6 participants who did not develop pharyngitis. Where serum IgG responses were produced, they were maintained for at least 3 months, which suggests that *S. pyogenes* pharyngeal infection can induce long-lived antibody secreting cells^52^, as serum antibodies would have waned substantially based on IgG serum half-life alone^53^. Overall, the post-challenge antibody responses described here add to recent longitudinal studies^12,15,51^ undermining the historical conception of asymptomatic colonisation (‘carriage’) with *S. pyogenes* as immunologically inert.

The mucosal and systemic pre-challenge antibody profiles of CHIVAS-M75 participants fits with repeated *S. pyogenes* exposures throughout early life, driving development of humoral immune memory. The inverse correlations we observed between baseline antibody levels and post-challenge responses resemble antibody feedback-mediated suppression of B cell responses, the basis for anti-D prophylaxis during pregnancy for Rhesus-negative women^54^ and recently reported to influence antibody responses to SARS-CoV-2 and malaria vaccines^55,56^ ^57^. This phenomenon is distinct from ‘original antigenic sin’^58^, which would have boosted rather than suppressed antibody responses from pre-existing B cell memory, particularly against the highly conserved antigens. Although it remains to be seen whether antibody feedback will affect *S. pyogenes* vaccines, it is reassuring that increased antigen dose can overcome feedback in mice^59^, suggesting that vaccine design may overcome suppression.

The limitations of the human *S. pyogenes* pharyngitis model have been discussed in depth previously^19,35,37^ and broadly relate to generalisability^34^, including age of participants, direct inoculation, and use of a single strain, among others. Reassuringly, we showed that pre-and post-challenge antibody levels among healthy and convalescent adults resembled those of healthy and convalescent children, the initial target group for vaccine development. The challenge strain, *emm*75, ranks among the top 10 most frequently isolated across all settings^60^. The confirmed inoculum from CHIVAS-M75 is at least 20 times lower than the dose delivered in 1970s human trials which successfully demonstrated vaccine-induced protection^17,18,35^ and up to 10,000 times lower than in animal models used in pre-clinical vaccine evaluation^25^. This argues against concerns the human model may be too stringent to support assessment of natural and vaccine-induced protective immunity. The major limitation of the immunological analyses is that the sample size, including just 6 participants without pharyngitis, precluded definitive conclusions regarding antigen-specific antibody results related to protection or susceptibility to pharyngitis. More fundamentally, whether our findings may contribute to defining a correlate of natural protection in future challenge and/or longitudinal surveillance studies is an open question, and its highly possible that correlates of protection from vaccination will be different.

Human challenge trials are poised to evaluate the efficacy of *S. pyogenes* vaccines for the first time in almost 50 years. Our findings highlight the power of the experimental human *S. pyogenes* pharyngitis model as a platform for investigating systemic and mucosal immunity. Clinical and immunological responses to *S. pyogenes* exposure are influenced by an individual’s imprint of past infections, underscoring the importance of assessing pre-existing immunity in planned vaccine trials in adults and children. This work will facilitate future studies that link responses to vaccination and infection with subsequent clinical outcomes, to inform interpretation of early phase vaccine trial results, and to converge on correlates of protection to support vaccine development.

## Methods

### Experimental pharyngitis model

Details of the CHIVAS-M75 study protocol, *emm*75 *S. pyogenes* challenge strain, and trial results have been described previously^19,35,37^. In the CHIVAS-M75 trial, 19 of 25 participants were diagnosed with pharyngitis, comprising 18/20 that received a dose of 10^5^ colony-forming units (CFU)/ml, and 1/5 that were given a lower dose (10^4^ CFU/ml). The clinical picture of acute pharyngitis was supported by microbiological (qPCR and culture), biochemical (C-reactive protein, cytokines and chemokines) and clinical streptococcal serology (anti-streptolysin O, anti-DnaseB) results. Considering all available data, one participant (SN057) who was not contemporaneously diagnosed with pharyngitis^35^ clearly did have pharyngitis and they have been categorised as such in this study. All participants with pharyngitis had *S. pyogenes* detected by throat swab qPCR and culture at multiple timepoints^61^. *S. pyogenes* colonisation was detected in only one of the 6 participants (participant SN013) who did not develop pharyngitis^35^.

### CHIVAS-M75 samples

Serum and saliva were collected from participants at screening, the evening prior to challenge, 8 to 12 hours after challenge, daily during the remainder of the inpatient period, then at 1-week, 1-month, 3-month, and 6-month outpatient visits. Blood was collected in serum separator tubes (BD Vacutainer SST Gold 8.5 mL) and allowed to clot. Within 2 hours of collection, tubes were centrifuged at 1500 x *g* for 15 minutes at 20°C, and then aliquots were stored at −80°C. Saliva samples were collected after a 30-minute fast from food and drink beginning with an initial brief mouth rinse with 50-100 mL of tap water, repeated after 25 minutes. After 30 minutes, participants began depositing fluid into a 50 mL Falcon tube standing upright in a cup of ice, to a total of 5-10 mL over the next 30 minutes. The fluid was then centrifuged at 2,600 x *g* for 15 minutes at 4°C, then phenylmethylsulfonyl fluoride (Sigma) was added to a final concentration of 500 nM, and aliquots were stored at −80°C until analysis.

### Antigens

A panel of 17 *S. pyogenes* vaccine candidate antigens, comprising recombinant proteins, synthetic peptides, and purified carbohydrate (Table S1) was investigated. Synthetic peptides from the M75 protein hypervariable region (M75-HVR), M-related protein (Mrp) 24, and Enn336 protein, were produced by GenScript with 95% purity, and collaborators at Griffith University provided p145, J8, P*17, and K4S2^62^. Recombinant proteins produced in BL21(DE3) pLysS *E. coli* representing the full length M75 protein^21^, T25 pilus, and TeeVax-3^63^ were provided by collaborators at the University of Auckland. University of Queensland collaborators provided recombinant proteins representing arginine deiminase (ADI), trigger factor (TF), and C5a peptidase (ScpA)^64^. Recombinant proteins representing *S. pyogenes* Adhesion and Division protein (SpyAD), Cell Envelope Protease (SpyCEP), and Streptolysin O (SLO), plus purified native Group A Carbohydrate (GAC)^65^, came from GSK Vaccines Institute for Global Health.

### Enzyme linked immunosorbent assays (ELISA)

ELISAs were done for all 17 antigens using serum and saliva from all 25 CHIVAS-M75 participants. In this study, data from ELISAs performed using samples from 2 time-points were analysed for serum IgG (pre-challenge and 1-month, or 3-month for 1 participant who failed to attend a 1-month follow up visit) and saliva IgA (pre-challenge and 1-week follow up visit). All antigens were resuspended to 5 µg/mL in 0.1 M coating carbonate buffer (0.03 M Na_2_CO_3_, 0.07 M NaHCO_3_, pH 9.6), except full-length M75 protein and GAC which were used at 1 µg/mL and coated at 50 µL per well onto 96-well medium-binding ELISA plates (Greiner Bio-One) overnight at room temperature. Plates were then blocked with 200 µL of 2% w/v bovine serum albumin (BSA, Sigma) in phosphate-buffered saline (PBS) for 1 or 2 hours at 37°C for IgG and IgA, respectively, followed by 5 washes in wash buffer (PBS with 0.05% v/v Tween20, PBS-T). Serial dilutions were then performed using diluent (PBS with 0.5% w/v BSA and 0.05% v/v Tween20) for sera (1:100, 1:300, 1:900, 1:2700) or saliva (1:50, 1:100, 1:200, 1:400) and then 50 µL added per well in duplicate. A set of healthy adult sera and saliva with high reactivity against the antigen panel were included as standardisation controls on each plate in duplicate; sera at 1:100, 1:200, 1:400, 1:800, 1:1600, 1:3200, 1:6400, and saliva at 1:50, 1:100, 1:200, 1:400, 1:800, 1:1600, 1:3200 dilutions. Plates were then incubated for one hour at 37°C (orbital shaking for saliva) then washed five times with PBS-T. Negative controls included both antigen-coated wells without serum and blocked wells with positive control serum, all of which had an absorbance at OD_450nm_ of approximately 0.05 across all plates. Horseradish peroxidase-labelled goat anti-human IgG (IgG-HRP, diluted 1:5000) or IgA (IgA-HRP, diluted 1:2000) secondary antibodies (Southern Biotech) were added at 50 µL/well for one hour at 37°C, followed by 5 washes with PBS-T and 3 washes with reverse osmosis water. Tetramethylbenzidine substrate (Life Technologies) was added at 50 µL/well and developed at room temperature for 1 minute (SLO, GAC, full-length M75 protein) or 5 minutes (all other antigens). Reactions were stopped with 1 M phosphoric acid and then absorbance was measured at OD_450nm_. Serum IgG end-point titres for each antigen and each sample are shown in Fig S4. Results were discarded and the assay repeated if the R^2^ of the reference curve of the positive standardisation control was ≤0.993 or the standard deviation between replicates was high. Human salivary IgA was quantified using the Human IgA SimpleStep ELISA® Kit (Abcam) according to the manufacturer’s instructions, with 1:5000 and 1:10000 dilutions tested.

### Multiplex bead assays

Antigen-specific IgG antibody responses were measured using a previously described multiplex bead-based assay (Luminex) ^43,44^. Briefly, beads coupled with *S. pyogenes* antigens were incubated with sera diluted 1:8000 in assay buffer (PBS pH 7.4, containing 1% IgG-free BSA). Following washing, bound IgG was detected with phycoerythrin labelled anti-human IgG (Jackson ImmunoResearch Labs). Median fluorescence intensity signals for each sample were measured with a MagPix instrument and the xPonent software version 4.2 (Luminex Corporation). Standard curves were generated from pooled human immunoglobulin (Privigen; CSL Behring), which was buffer exchanged into PBS, and fitted using 5-parameter logistic regression with Belysa software (Merck). Antigen-specific IgG concentrations in μg/mL were quantified by interpolation.

Paediatric comparator data are from a previously published study of *S. pyogenes* infections from Auckland, New Zealand^44,46^. Available data are shown for children (5-14 years old) with *S. pyogenes* pharyngitis (n=24) and healthy controls (n=15) who were deemed low risk for developing serious *S. pyogenes* disease, including acute rheumatic fever. Convalescent sera were obtained approximately 4 weeks after microbiologically-confirmed *S. pyogenes* positive pharyngitis and were included in the comparator analysis if a serological response to at least one antigen in the multiplex bead-based assay was observed when compared to matched healthy children. For both datasets, only data for the 6 relevant antigens were analysed for this study.

### Bacterial isolates and growth

M75 *Streptococcus pyogenes* bacteria (611024, accession number CP033621)^37^ were grown stationary at 37℃ with 5% CO_2_ in Todd-Hewitt broth (Oxoid) with 1% (w/v) yeast extract (THY; Bacto) or on solid THY-agar with 0.005% 2,3,5-tetraphenyltetrazolium chloride (TTC; Sigma).

### Opsonophagocytic killing assays (OPKA)

OPKAs were carried out as previously described^21,38,66^. Briefly, HL-60 cells were differentiated for 3 or 4 days using 0.8% dimethylformamide at 37°C with 5% CO_2_ and diluted in opsonization buffer (10% v/v heat-inactivated fetal bovine serum (HyClone)), 0.1% w/v gelatin (Sigma) in Hanks’ balanced salt solution with Ca/Mg) to 1 x 10^7^ cells/ml. Frozen *emm*75 bacterial stocks were thawed, washed, and diluted in opsonization buffer to ∼120,000 CFU/ml and pre-incubated for 30 minutes at room temperature with 20 µl heat-inactivated participant sera. Baby rabbit complement and differentiated HL-60 cells were then added and incubated for 60 minutes at 37°C with 5% CO_2_. Surviving bacteria were recovered, then enumerated by dilution on TTC-THY agar as previously described^21^. Bacteria were enumerated using a ProtoCOL3 automated colony counter (Synbiosis). The dilution of sera resulting in 50% killing was calculated as the opsonic index using the Opsotiter software (license obtained from the Pathogen Reference Laboratory at the University of Alabama at Birmingham^67^). Positive and negative control sera was included in every assay.

### Bacterial adhesion assays

The adhesion assay using Detroit 562 (D562) human pharyngeal cells was performed as described previously^37,39^, with the addition of a pre-incubation step with saliva described below. D562 cells were maintained in assay medium (Dulbecco’s Modified Eagle Medium with 10% fetal bovine serum, and 1x Pen-Strep (Sigma)). M75 *S. pyogenes* was grown to mid-exponential phase (OD_600nm_ ∼0.5), washed, and resuspended in 1 ml of assay media. Input inoculum per test was ∼10^6^ CFU/well to achieve a multiplicity of infection of 5 *S. pyogenes* CFU to one D562 cell. Bacteria were pre-incubated for 30 minutes at 37℃ with participant saliva (40 µl) collected pre-challenge and 1-week following discharge, then added to D562 cells plated in 360 µL of assay medium per well of 24-well tissue culture plates (Nunc) in triplicate. Plates were then centrifuged for 5 mins at 200 ×*g* before incubation for 1 hour at 37°C in 5% CO_2_. After the incubation, non-adherent bacteria were removed by washing three times with 1 mL PBS, then total cell-associated bacteria (adherent plus invasive) were recovered by adding 200 µl 0.25% trypsin, then lysis with 0.025% Triton-X (Sigma) in distilled water. Bacteria were then enumerated by track dilution on TTC-THY agar as previously described^68^. M75 *S. pyogenes* adhesion (%) was determined as CFU in saliva sample wells relative to control wells (without saliva). Triplicate wells were averaged, and assays performed at least twice per sample.

### Statistical analysis

Analysis of raw data and data extrapolation was performed using Prism (GraphPad). Statistical analysis was conducted in R version 3.8^69^, using packages ‘ggplot2, ‘pheatmap’, ‘viridis’, ‘tidyverse’, ‘reshape2’, ‘ggpubr’, ‘RColorBrewer’, ‘gghalves’, ‘vegan’, ‘ggbeeswarm’, ‘mixOmics’, ‘Polychrome’, and ‘uwot’. For serum IgG ELISAs, a four-parameter logistic curve was generated from the standard curve for each antigen for each plate, and a titre for each serum sample was calculated relative to that curve and expressed as arbitrary units (AU). For antigen-specific saliva IgA ELISAs, sample AU values were calculated relative to the reference control saliva at the 1:50 dilution. Participants were deemed to have a ‘response’ where the AU post-challenge was 25% greater than the pre-challenge value. This cut off was used to determine: 1) the proportion of individuals who responded to each antigen; and 2) the number of antigens to which each participant responded. For total IgA, values were extrapolated from a log-transformed 5-parametric logistical standard curve, generated using known amounts of IgA. P-values were determined using non-parametric methods (Mann-Whitney tests for unpaired and Wilcoxon-signed rank test for paired 2 group comparisons). Where repeated 2 group comparisons were performed across the panel of antigens, p-values were adjusted for multiple corrections using the false discovery rate (FDR) approach ^70^. For 3 or more groups, paired sample comparisons were performed using Friedman’s test followed by Wilcoxon signed-rank test with FDR-adjustments, and between group comparisons using Kruskall-Wallis and Dunn’s multiple comparison testing.

Multidimensional scaling was performed using the ‘vegan’ package using ELISA or multiplex bead-based assay data for 18 antigens that were each z-score transformed. A matrix of Spearman Correlation coefficients, comparing each participant’s antibody responses to every other participant, was generated and used to calculate a distance matrix for each person and each variable. This was used to perform non-metric multidimensional scaling to reduce the dimensionality of the data to 2, as previously described^71^. The coordinates for Dimension 1 from the multidimensional scaling analysis were compared between groups, as well as subtracting pre-challenge from 1-month coordinates.

Sparse partial least squares discriminant analysis was performed using the mixOmics package. ELISA variables were used except where multiplex bead-based assay data was available. Correlation was determined by Spearman correlation coefficient for clinical variables on discrete numerical scales, described previously^35^. The initial model allowed for 10 components, model performance evaluated down to 2 components, then tuned using 3 Mfold validation and 100 iterations which returned 5 immune variables and 1 dimension as optimal to explain the difference between pharyngitis and non-pharyngitis participants.

## Acknowledgements

We thank Prof. Thomas Proft and Dr Jacelyn Loh (University of Auckland) for providing T25 and TeeVax-3 antigens. We thank Dr Danilo Gomes Moriel (GSK Vaccines Institute for Global Health) for supplying GAC, SLO, SpyCEP, and SpyAD antigens, including for development of the Luminex multiplex bead-based assay. GSK Vaccines Institute for Global Health is an affiliate of GlaxoSmithKline Biologicals SA. We are grateful to Dr Julie Bennett and Prof. Michael Baker at the University of Otago, Wellington, New Zealand, and all contributors to the study from which comparator IgG data was obtained for streptococcal pharyngitis in children. Figures 1A, 2A and 3A were created with BioRender.com.

## References

1. Brouwer S, Rivera-Hernandez T, Curren BF, et al. Pathogenesis, epidemiology and control of Group A Streptococcus infection. Nat Rev Microbiol 2023;21(7):431–447. DOI: 10.1038/s41579-023-00865-7.

2. Roth GA, Mensah GA, Johnson CO, et al. Global Burden of Cardiovascular Diseases and Risk Factors, 1990-2019: Update From the GBD 2019 Study. J Am Coll Cardiol 2020;76(25):2982–3021. DOI: 10.1016/j.jacc.2020.11.010.

3. Ralph AP, Carapetis JR. Group a streptococcal diseases and their global burden. Curr Top Microbiol Immunol 2013;368:1–27. DOI: 10.1007/82_2012_280.

4. Immunization VaB, Vaccine Prioritization & Platforms Team. Strategic Priority 7: Regional engagement strategy to identify IA2030 global priority endemic pathogens. Product Development for Vaccines Advisory Committee (PDVAC) Meeting, World Health Organization. (https://www.who.int/news-room/events/detail/2023/12/12/default-calendar/2023-meeting-who-product-development-for-vaccines-advisory-committee-meeting-pdvac).

5. Osowicki J, Vekemans J, Kaslow DC, Friede MH, Kim JH, Steer AC. WHO/IVI global stakeholder consultation on group A Streptococcus vaccine development: Report from a meeting held on 12-13 December 2016. Vaccine 2018;36(24):3397–3405. DOI: 10.1016/j.vaccine.2018.02.068.

6. Vekemans J, Gouvea-Reis F, Kim JH, et al. The Path to Group A Streptococcus Vaccines: World Health Organization Research and Development Technology Roadmap and Preferred Product Characteristics. Clin Infect Dis 2019;69(5):877–883. DOI: 10.1093/cid/ciy1143.

7. Walkinshaw DR, Wright MEE, Mullin AE, Excler JL, Kim JH, Steer AC. The Streptococcus pyogenes vaccine landscape. NPJ Vaccines 2023;8(1):16. DOI: 10.1038/s41541-023-00609-x.

8. Frost H, Excler JL, Sriskandan S, Fulurija A. Correlates of immunity to Group A Streptococcus: a pathway to vaccine development. NPJ Vaccines 2023;8(1):1. DOI: 10.1038/s41541-022-00593-8.

9. Lorenz N, Ho TKC, McGregor R, et al. Serological Profiling of Group A Streptococcus Infections in Acute Rheumatic Fever. Clin Infect Dis 2021;73(12):2322–2325. (In eng). DOI: 10.1093/cid/ciab180.

10. Tsoi SK, Smeesters PR, Frost HR, Licciardi P, Steer AC. Correlates of Protection for M Protein-Based Vaccines against Group A Streptococcus. J Immunol Res 2015;2015:167089. DOI: 10.1155/2015/167089.

11. Lancefield RC. Current knowledge of type-specific M antigens of group A streptococci. J Immunol 1962;89:307–13. (https://www.ncbi.nlm.nih.gov/pubmed/14461914).

12. Hysmith ND, Kaplan EL, Cleary PP, Johnson DR, Penfound TA, Dale JB. Prospective Longitudinal Analysis of Immune Responses in Pediatric Subjects After Pharyngeal Acquisition of Group A Streptococci. J Pediatric Infect Dis Soc 2017;6(2):187–196. DOI: 10.1093/jpids/piw070.

13. Guirguis N, Fraser DW, Facklam RR, El Kholy A, Wannamaker LW. Type-specific immunity and pharyngeal acquisition of group A Streptococcus. Am J Epidemiol 1982;116(6):933–9. (In eng). DOI: 10.1093/oxfordjournals.aje.a113495.

14. Wannamaker LW, Denny FW, Perry WD, Siegel AC, Rammelkamp CH, Jr. Studies on immunity to streptococcal infections in man. AMA Am J Dis Child 1953;86(3):347–8. (In eng).

15. Salie MT, Muhamed B, Engel K, et al. Serum Immune Responses to Group A Streptococcal Antigens following Pharyngeal Acquisitions among Children in Cape Town, South Africa. mSphere 2023;8(3):e00113–23. DOI: 10.1128/msphere.00113-23.

16. D’Alessandri R, Plotkin G, Kluge RM, et al. Protective studies with group A streptococcal M protein vaccine. III. Challenge of volunteers after systemic or intranasal immunization with Type 3 or Type 12 group A Streptococcus. J Infect Dis 1978;138(6):712–8. (In eng) (https://academic.oup.com/jid/article-abstract/138/6/712/790110?redirectedFrom=fulltext).

17. Fox EN, Waldman RH, Wittner MK, Mauceri AA, Dorfman A. Protective study with a group A streptococcal M protein vaccine. Infectivity challenge of human volunteers. J Clin Invest 1973;52(8):1885–92. (In eng). DOI: 10.1172/jci107372.

18. Polly SM, Waldman RH, High P, Wittner MK, Dorfman A. Protective studies with a group A streptococcal M protein vaccine. II. Challenge of volunteers after local immunization in the upper respiratory tract. J Infect Dis 1975;131(3):217–24. (In eng).

19. Osowicki J, Azzopardi KI, Baker C, et al. Controlled human infection for vaccination against Streptococcus pyogenes (CHIVAS): Establishing a group A Streptococcus pharyngitis human infection study. Vaccine 2019;37(26):3485–3494. (In eng). DOI: 10.1016/j.vaccine.2019.03.059.

20. Rivera-Hernandez T, Rhyme MS, Cork AJ, et al. Vaccine-Induced Th1-Type Response Protects against Invasive Group A Streptococcus Infection in the Absence of Opsonizing Antibodies. mBio 2020;11(2). DOI: 10.1128/mBio.00122-20.

21. McGregor R, Paterson A, Sharma P, et al. Naturally acquired functional antibody responses to group A Streptococcus differ between major strain types. mSphere 2023;8(5):e0017923. (In eng). DOI: 10.1128/msphere.00179-23.

22. Gill FA. A review of past attempts and present concepts of producing streptococcal immunity in humans. Q Bull Northwest Univ Med Sch 1960;34:326–39. (https://www.ncbi.nlm.nih.gov/pubmed/13705305).

23. Revocation of status of specific products; Group A streptococcus. Direct final rule. Fed Regist 2005;70(231):72197–9. (In eng).

24. Asturias EJ, Excler JL, Ackland J, et al. Safety of Streptococcus pyogenes Vaccines: Anticipating and Overcoming Challenges for Clinical Trials and Post-Marketing Monitoring. Clin Infect Dis 2023;77(6):917–924. (In eng). DOI: 10.1093/cid/ciad311.

25. Harbison-Price N, Rivera-Hernandez T, Osowicki J, et al. Current Approaches to Vaccine Development of Streptococcus pyogenes. In: Ferretti JJ, Stevens DL, Fischetti VA, eds. Streptococcus pyogenes: Basic Biology to Clinical Manifestations. Oklahoma City (OK): University of Oklahoma Health Sciences Center © The University of Oklahoma Health Sciences Center.; 2022.

26. Reglinski M, Gierula M, Lynskey NN, Edwards RJ, Sriskandan S. Identification of the Streptococcus pyogenes surface antigens recognised by pooled human immunoglobulin. Sci Rep 2015;5:15825. (In eng). DOI: 10.1038/srep15825.

27. Bensi G, Mora M, Tuscano G, et al. Multi high-throughput approach for highly selective identification of vaccine candidates: the Group A Streptococcus case. Molecular & cellular proteomics : MCP 2012;11(6):M111.015693. (In eng). DOI: 10.1074/mcp.M111.015693.

28. Sekuloski S, Batzloff MR, Griffin P, et al. Evaluation of safety and immunogenicity of a group A streptococcus vaccine candidate (MJ8VAX) in a randomized clinical trial. PloS one 2018;13(7):e0198658. (In eng). DOI: 10.1371/journal.pone.0198658.

29. Pastural E, McNeil SA, MacKinnon-Cameron D, et al. Safety and immunogenicity of a 30-valent M protein-based group a streptococcal vaccine in healthy adult volunteers: A randomized, controlled phase I study. Vaccine 2020;38(6):1384–1392. DOI: 10.1016/j.vaccine.2019.12.005.

30. McNeil SA, Halperin SA, Langley JM, et al. A double-blind, randomized phase II trial of the safety and immunogenicity of 26-valent group A streptococcus vaccine in healthy adults. International Congress Series 2006;1289:303–306. DOI: 10.1016/j.ics.2005.12.002.

31. McNeil SA, Halperin SA, Langley JM, et al. Safety and immunogenicity of 26-valent group A streptococcus vaccine in healthy adult volunteers. Clinical Infectious Diseases 2005;41(8):1114–1122. (In English). DOI: 10.1086/444458.

32. Kotloff KL, Corretti M, Palmer K, et al. Safety and immunogenicity of a recombinant multivalent group A streptococcal vaccine in healthy adults: Phase 1 trial. Journal of the American Medical Association 2004;292(6):709–715. (In English). DOI: 10.1001/jama.292.6.709.

33. Steer AC, Carapetis JR, Dale JB, et al. Status of research and development of vaccines for Streptococcus pyogenes. Vaccine 2016;34(26):2953–8. (In eng). DOI: 10.1016/j.vaccine.2016.03.073.

34. Abo YN, Jamrozik E, McCarthy JS, Roestenberg M, Steer AC, Osowicki J. Strategic and scientific contributions of human challenge trials for vaccine development: facts versus fantasy. Lancet Infect Dis 2023;23(12):e533–e546. DOI: 10.1016/S1473-3099(23)00294-3.

35. Osowicki J, Azzopardi KI, Fabri L, et al. A controlled human infection model of Streptococcus pyogenes pharyngitis (CHIVAS-M75): an observational, dose-finding study. Lancet Microbe 2021;2(7):e291–e299. DOI: 10.1016/S2666-5247(20)30240-8.

36. Anderson J, Imran S, Frost HR, et al. Immune signature of acute pharyngitis in a Streptococcus pyogenes human challenge trial. Nat Commun 2022;13(1):769. (In eng). DOI: 10.1038/s41467-022-28335-3.

37. Osowicki J, Azzopardi KI, McIntyre L, et al. A Controlled Human Infection Model of Group A Streptococcus Pharyngitis: Which Strain and Why? mSphere 2019;4(1). DOI: 10.1128/mSphere.00647-18.

38. McGregor R, Jones S, Jeremy RM, Goldblatt D, Moreland NJ. An Opsonophagocytic Killing Assay for the Evaluation of Group A Streptococcus Vaccine Antisera. Methods Mol Biol 2020;2136:323–335. (In eng). DOI: 10.1007/978-1-0716-0467-0_26.

39. Khemlani AHJ, Proft T, Loh JMS. Assays to Analyze Adhesion of Group A Streptococcus to Host Cells. Methods Mol Biol 2020;2136:271–278. (In eng). DOI: 10.1007/978-1-0716-0467-0_20.

40. Collin M, Olsén A. Effect of SpeB and EndoS from Streptococcus pyogenes on human immunoglobulins. Infect Immun 2001;69(11):7187–9. (In eng). DOI: 10.1128/iai.69.11.7187-7189.2001.

41. Zhu H, Chelysheva I, Cross DL, et al. Molecular correlates of vaccine-induced protection against typhoid fever. The Journal of clinical investigation 2023;133(16). DOI: 10.1172/JCI169676.

42. Knuutila A, Barkoff AM, Mertsola J, Osicka R, Sebo P, He Q. Simultaneous Determination of Antibodies to Pertussis Toxin and Adenylate Cyclase Toxin Improves Serological Diagnosis of Pertussis. Diagnostics (Basel) 2021;11(2). DOI: 10.3390/diagnostics11020180.

43. Whitcombe AL, Han F, McAlister SM, et al. An eight-plex immunoassay for Group A streptococcus serology and vaccine development. J Immunol Methods 2022;500:113194. DOI: 10.1016/j.jim.2021.113194.

44. Whitcombe AL, McGregor R, Bennett J, et al. Increased Breadth of Group A Streptococcus Antibody Responses in Children With Acute Rheumatic Fever Compared to Precursor Pharyngitis and Skin Infections. J Infect Dis 2022;226(1):167–176. DOI: 10.1093/infdis/jiac043.

45. Hanson-Manful P, Whitcombe AL, Young PG, et al. The novel Group A Streptococcus antigen SpnA combined with bead-based immunoassay technology improves streptococcal serology for the diagnosis of acute rheumatic fever. J Infect 2018;76(4):361–368. (In eng). DOI: 10.1016/j.jinf.2017.12.008.

46. Bennett J, Moreland NJ, Zhang J, et al. Risk factors for group A streptococcal pharyngitis and skin infections: A case control study. Lancet Reg Health West Pac 2022;26:100507. DOI: 10.1016/j.lanwpc.2022.100507.

47. Henningham A, Chiarot E, Gillen CM, et al. Conserved anchorless surface proteins as group A streptococcal vaccine candidates. J Mol Med (Berl) 2012;90(10):1197–207. (In eng). DOI: 10.1007/s00109-012-0897-9.

48. Ozberk V, Pandey M, Good MF. Contribution of cryptic epitopes in designing a group A streptococcal vaccine. Human Vaccines & Immunotherapeutics 2018;14(8):2034–2052. DOI: 10.1080/21645515.2018.1462427.

49. Carducci M, Whitcombe A, Rovetini L, et al. Development and characterization of a hemolysis inhibition assay to determine functionality of anti-Streptolysin O antibodies in human sera. J Immunol Methods 2024;526:113618. DOI: 10.1016/j.jim.2024.113618.

50. Turner CE, Kurupati P, Wiles S, Edwards RJ, Sriskandan S. Impact of immunization against SpyCEP during invasive disease with two streptococcal species: Streptococcus pyogenes and Streptococcus equi. Vaccine 2009;27(36):4923–9. DOI: 10.1016/j.vaccine.2009.06.042.

51. Keeley AJ, Groves D, Armitage EP, et al. Streptococcus pyogenes Colonization in Children Aged 24–59 Months in the Gambia: Impact of Live Attenuated Influenza Vaccine and Associated Serological Responses. The Journal of Infectious Diseases 2023;228(7):957–965. DOI: 10.1093/infdis/jiad153.

52. Corcoran LM, Tarlinton DM. Regulation of germinal center responses, memory B cells and plasma cell formation-an update. Curr Opin Immunol 2016;39:59–67. (In eng). DOI: 10.1016/j.coi.2015.12.008.

53. Mankarious S, Lee M, Fischer S, et al. The half-lives of IgG subclasses and specific antibodies in patients with primary immunodeficiency who are receiving intravenously administered immunoglobulin. J Lab Clin Med 1988;112(5):634–40. (In eng).

54. Bowman JM. The prevention of Rh immunization. Transfus Med Rev 1988;2(3):129–50. (In eng). DOI: 10.1016/s0887-7963(88)70039-5.

55. Schaefer-Babajew D, Wang Z, Muecksch F, et al. Antibody feedback regulates immune memory after SARS-CoV-2 mRNA vaccination. Nature 2023;613(7945):735–742. (In eng). DOI: 10.1038/s41586-022-05609-w.

56. Inoue T, Shinnakasu R, Kawai C, et al. Antibody feedback contributes to facilitating the development of Omicron-reactive memory B cells in SARS-CoV-2 mRNA vaccinees. Journal of Experimental Medicine 2022;220(2). DOI: 10.1084/jem.20221786.

57. McNamara HA, Idris AH, Sutton HJ, et al. Antibody Feedback Limits the Expansion of B Cell Responses to Malaria Vaccination but Drives Diversification of the Humoral Response. Cell host & microbe 2020;28(4):572–585.e7. (In eng). DOI: 10.1016/j.chom.2020.07.001.

58. Koutsakos M, Ellebedy AH. Immunological imprinting: Understanding COVID-19. Immunity 2023;56(5):909–913. (In eng). DOI: 10.1016/j.immuni.2023.04.012.

59. Tas JMJ, Koo JH, Lin YC, et al. Antibodies from primary humoral responses modulate the recruitment of naive B cells during secondary responses. Immunity 2022;55(10):1856–1871.e6. (In eng). DOI: 10.1016/j.immuni.2022.07.020.

60. Smeesters PR, de Crombrugghe G, Tsoi SK, et al. Global Streptococcus pyogenes strain diversity, disease associations, and implications for vaccine development: a systematic review. The Lancet Microbe 2024;5(2):e181–e193. DOI: 10.1016/S2666-5247(23)00318-X.

61. Fabri LV, Azzopardi KI, Osowicki J, Frost HR, Smeesters PR, Steer AC. An emm-type specific qPCR to track bacterial load during experimental human Streptococcus pyogenes pharyngitis. BMC Infectious Diseases 2021;21(1):463. DOI: 10.1186/s12879-021-06173-w.

62. Pandey M, Mortensen R, Calcutt A, et al. Combinatorial Synthetic Peptide Vaccine Strategy Protects against Hypervirulent CovR/S Mutant Streptococci. The Journal of Immunology 2016;196(8):3364–3374. DOI: 10.4049/jimmunol.1501994.

63. Loh JMS, Rivera-Hernandez T, McGregor R, et al. A multivalent T-antigen-based vaccine for Group A Streptococcus. Sci Rep 2021;11(1):4353. (In eng). DOI: 10.1038/s41598-021-83673-4.

64. Rivera-Hernandez T, Carnathan DG, Jones S, et al. An Experimental Group A Streptococcus Vaccine That Reduces Pharyngitis and Tonsillitis in a Nonhuman Primate Model. MBio 2019;10(2) (In eng). DOI: 10.1128/mBio.00693-19.

65. Di Benedetto R, Mancini F, Carducci M, et al. Rational Design of a Glycoconjugate Vaccine against Group A Streptococcus. International Journal of Molecular Sciences 2020;21(22):8558. (https://www.mdpi.com/1422-0067/21/22/8558).

66. Jones S, Moreland NJ, Zancolli M, et al. Development of an opsonophagocytic killing assay for group a streptococcus. Vaccine 2018;36(26):3756–3763. (In eng). DOI: 10.1016/j.vaccine.2018.05.056.

67. Burton RL, Nahm MH. Development and validation of a fourfold multiplexed opsonization assay (MOPA4) for pneumococcal antibodies. Clin Vaccine Immunol 2006;13(9):1004–9. DOI: 10.1128/CVI.00112-06.

68. Frost HR, Tsoi SK, Baker CA, et al. Validation of an automated colony counting system for group A Streptococcus. BMC Res Notes 2016;9:72. DOI: 10.1186/s13104-016-1875-z.

69. R Core Team. R: A language and environment for statistical computing. Vienna, Austria: R Foundation for Statistical Computing; 2020.

70. Benjamini Y, Hochberg Y. Controlling the False Discovery Rate: A Practical and Powerful Approach to Multiple Testing. Journal of the Royal Statistical Society: Series B (Methodological) 1995;57(1):289–300. (https://rss.onlinelibrary.wiley.com/doi/abs/10.1111/j.2517-6161.1995.tb02031.x).

71. Hill DL, Carr EJ, Rutishauser T, et al. Immune system development varies according to age, location, and anemia in African children. Sci Transl Med 2020;12(529) (In eng). DOI: 10.1126/scitranslmed.aaw9522.

