## Supplementary Appendix for "*Streptococcus pyogenes* pharyngitis elicits diverse antibody responses to key vaccine antigens influenced by the imprint of past infections"

- **Table S1. Antigen panel**
- **Figure S1. Total saliva IgA before and after experimental challenge with *emm*75 *Streptococcus pyogenes.***
- **Figure S2. Antigen-specific serum IgG responses (ELISA) before and after *emm*75 *Streptococcus pyogenes* challenge**
- **Figure S3. Antigen-specific saliva IgA responses (ELISA) before and after *emm*75 *Streptococcus pyogenes* challenge**
- **Figure S4. ELISA end-point titres for 17 *Streptococcus pyogenes* antigens**

**Table S1. Antigen panel**

| Antigen | Description | Reference(s) |
|---|---|---|
| SpyCEP^ | Recombinant inactivated serine protease cleaving interleukin-8, impeding neutrophil chemotaxis | 1, 2 |
| SLO^ | Recombinant inactivated streptolysin O, a secreted, pore forming cytolysin | 2, 3, 4 |
| ScpA^ | Recombinant C5a peptidase, a protease cleaving C5a, interfering with neutrophil chemotaxis | 4, 5, 6, 7 |
| GAC^ | Purified group A carbohydrate | 3, 8, 9, 10 |
| M75 protein# | Full-length M75 recombinant protein | 11 |
| SpyAD^ | Recombinant protein, involved in adhesion to host cells and bacterial cell division | 2, 3, 12 |
| Mrp24# | Peptide from M-related protein | 13, 14, 15, 16 |
| Enn336# | Peptide from Enn protein | 15, 16 |
| T25# | T-antigen (pilus) | 17, 18 |
| TF^ | Trigger factor, recombinant anchorless surface protein contributing to protease secretion and activation | 19 |
| ADI^ | Arginine deiminase, recombinant anchorless surface protein involved in enzymatic conversion of arginine to citrulline | 19, 20 |
| M75 HVR# | First 50 amino acids of the M75 protein (N-terminal hypervariable region) | 21, 22, 23, 24 |
| TeeVax-3# (TV3.5d) | Multivalent recombinant protein joining N- and C-terminal domains of T-antigens (including T25) | 17, 18 |
| P*17^ | Synthetic 12 amino acid minimal B-cell epitope engineered from p145 | 25 |
| K4S2^ | 20 amino acid peptide B-cell epitope from SpyCEP made more soluble with four added lysine residues | 26 |
| p145^ | 20 amino acid peptide epitope from the M-protein C-repeat region | 27 |
| J8^ | 12 amino acid minimal B-cell epitope of p145 with flanking sequences | 28, 29 |
| SpnA^ | Recombinant *S. pyogenes* nuclease A, a cell wall-anchored DNA degrading enzyme | 30, 31 |
| DNAseB^ | Recombinant deoxyribonuclease-B/Streptodornase-B, a secreted DNA degrading enzyme | 30, 31 |

### Type-specific antigens (present in the *emm*75 *S. pyogenes* challenge strain)

^ Conserved or semi-conserved antigens

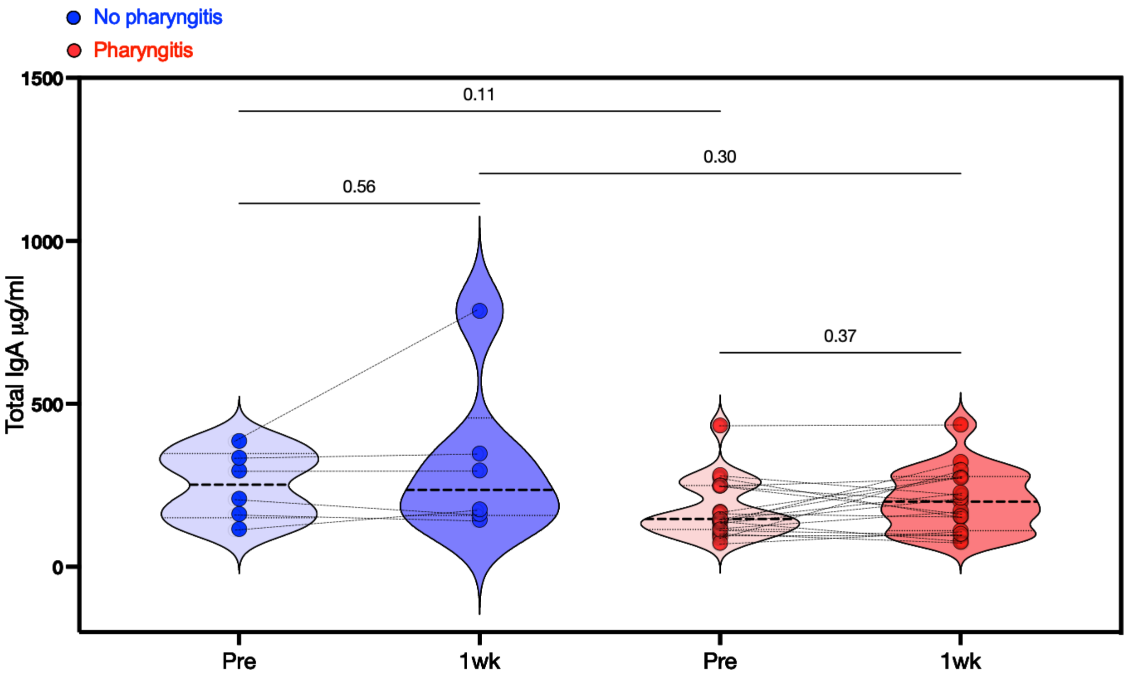


**Figure S1. Total saliva IgA before and after experimental challenge with *emm*75 *Streptococcus pyogenes.***

Quantification of total IgA present in saliva collected pre-challenge and 1-week after pharyngitis diagnosis (n=19) or discharge without pharyngitis (n=6). IgA concentration for each group shown as median (dotted line) and range with kernel density estimation, with within group comparisons performed using paired Wilcoxon signed-rank tests and between group comparisons using Mann-Whitney tests.

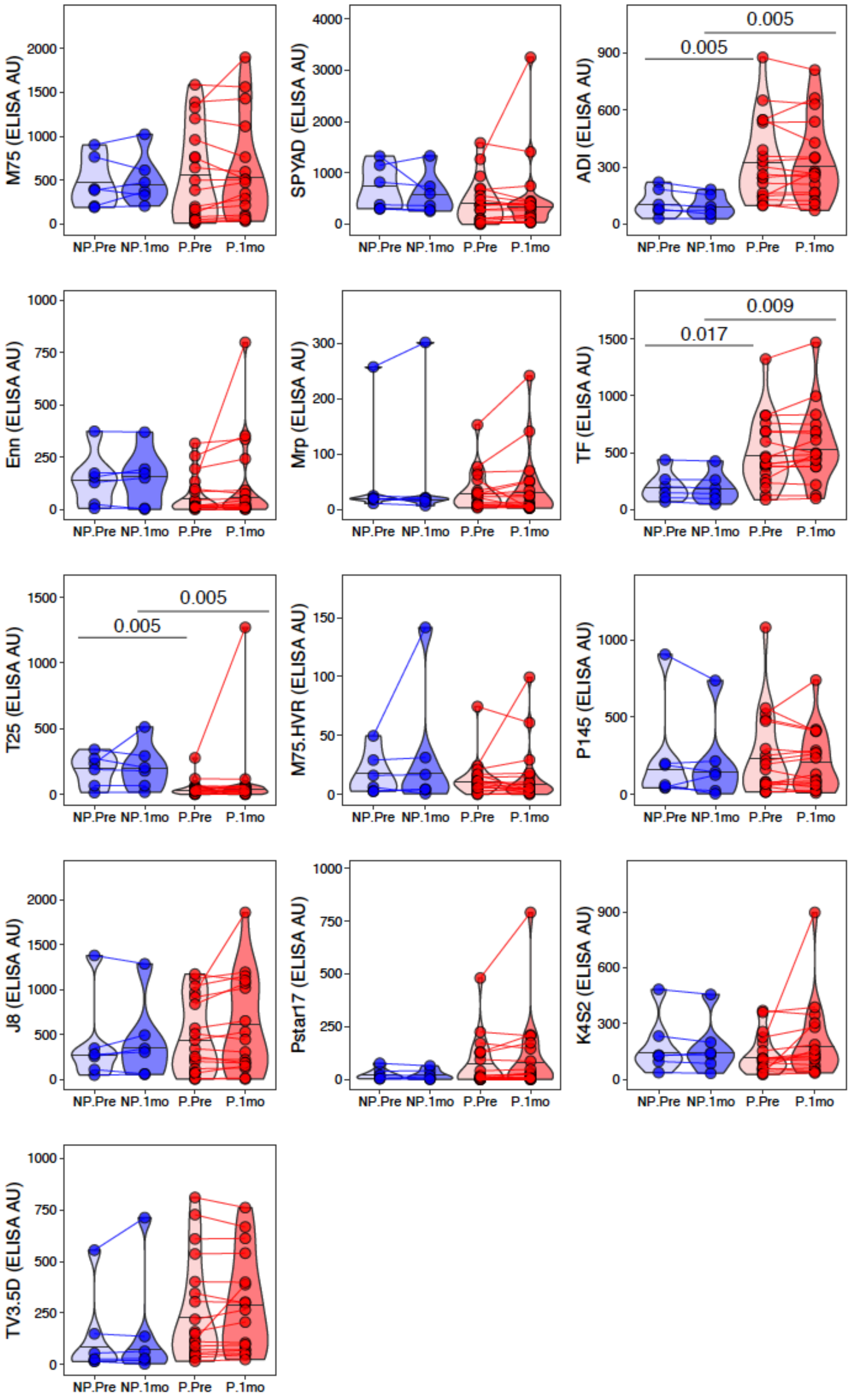


**Figure S2. Antigen-specific serum IgG responses (ELISA) before and after *emm*75 *Streptococcus pyogenes* challenge**

IgG arbitrary units (AU) for 13 antigens from pre and post challenge serum samples, split by pharyngitis outcome. Within group comparisons performed using paired Wilcoxon signed-rank test, and groups compared using Mann-Whitney test, all FDR-adjusted for multiple comparisons (pharyngitis n=19, no pharyngitis n=6).

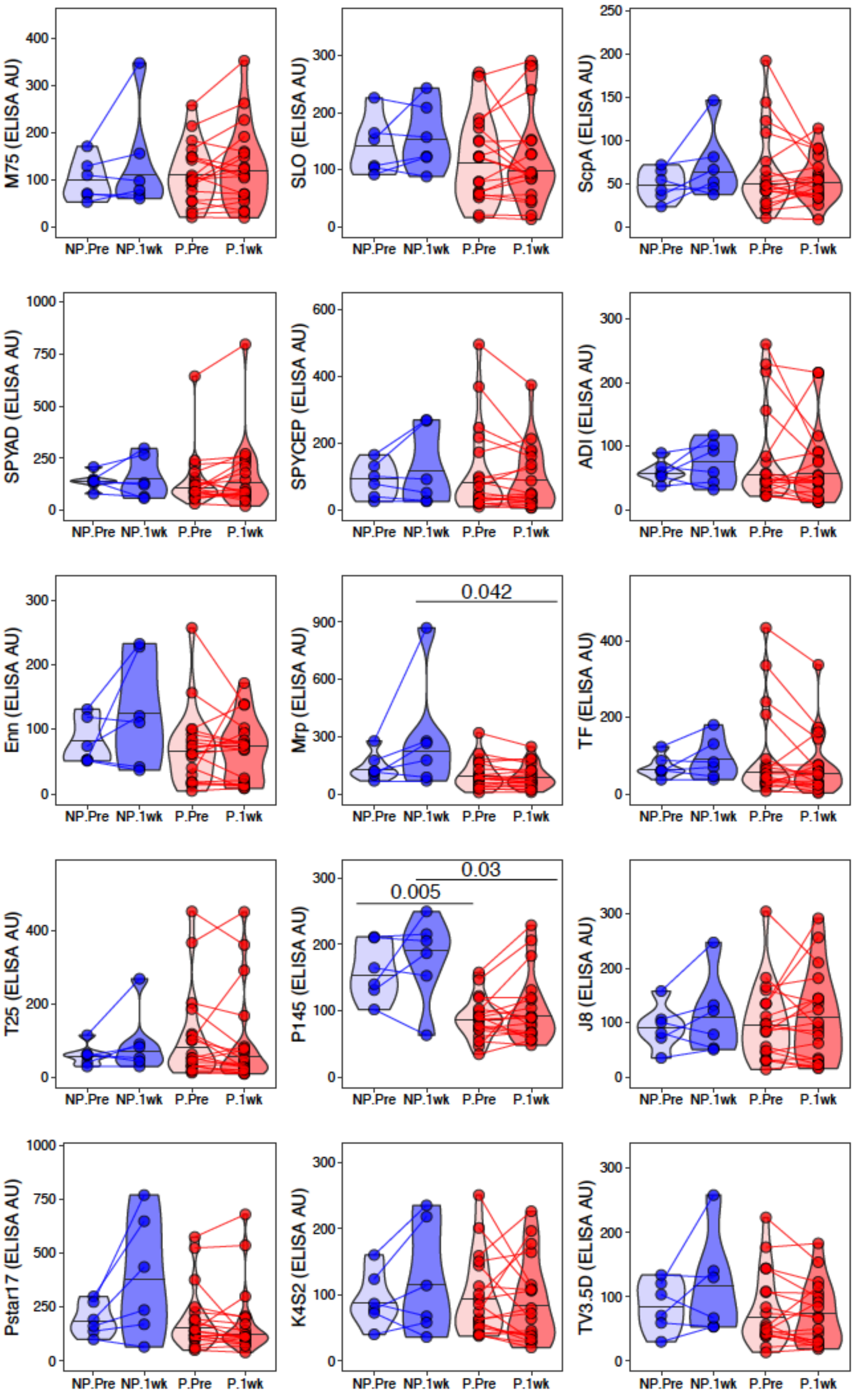


**Figure S3. Antigen-specific saliva IgA responses (ELISA) before and after *emm*75 *Streptococcus pyogenes* challenge.**

IgA arbitrary units (AU) for 15 antigens from pre and post challenge saliva samples, split by pharyngitis outcome. Within group comparisons performed using paired Wilcoxon signed-rank test, and groups compared using Mann-Whitney test, all FDR-adjusted for multiple comparisons (pharyngitis n=19, no pharyngitis n=6).

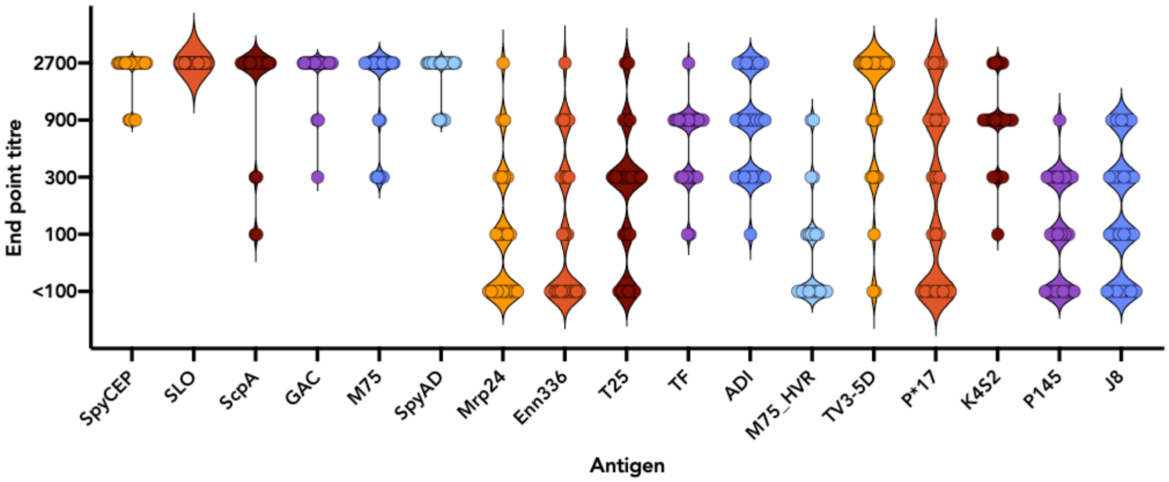


**Figure S4. ELISA end-point titres for 17 *Streptococcus pyogenes* antigens**

End-point titres calculated as the reciprocal of the highest dilution of serum with an $OD_{450nm}$ twice the value of the assay background. Each violin represents the distribution of 48 antibody titres from 1-month post-challenge for 24 CHIVAS participants, with individual titres represented by coloured circles.

#### References


1. Zingaretti, C. *et al.* Streptococcus pyogenes SpyCEP: a chemokine-inactivating protease with unique structural and biochemical features. *FASEB J* **24**, 2839-2848 (2010).

2. Bensi, G. *et al.* Multi high-throughput approach for highly selective identification of vaccine candidates: the Group A Streptococcus case. *Mol Cell Proteomics* **11**, M111 015693 (2012).

3. Di Benedetto, R. *et al.* Rational Design of a Glycoconjugate Vaccine against Group A Streptococcus. *Int J Mol Sci* **21** (2020).

4. Rivera-Hernandez, T. *et al.* Differing Efficacies of Lead Group A Streptococcal Vaccine Candidates and Full-Length M Protein in Cutaneous and Invasive Disease Models. *mBio* **7** (2016).

5. Cleary, P.P., Matsuka, Y.V., Huynh, T., Lam, H. & Olmsted, S.B. Immunization with C5a peptidase from either group A or B streptococci enhances clearance of group A streptococci from intranasally infected mice. *Vaccine* **22**, 4332-4341 (2004).

6. Park, H.-S. & Cleary, P.P. Active and passive intranasal immunizations with streptococcal surface protein C5a peptidase prevent infection of murine nasal mucosa-associated lymphoid tissue, a functional homologue of human tonsils. *Infect. Immun.* **73**, 7878-7886 (2005).

7. Shet, A., Kaplan, E.L., Johnson, D.R. & Cleary, P.P. Immune response to group A streptococcal C5a peptidase in children: implications for vaccine development. *J Infect Dis* **188**, 809-817 (2003).

8. Sabharwal, H. *et al.* Group A streptococcus (GAS) carbohydrate as an immunogen for protection against GAS infection. *J Infect Dis* **193**, 129-135 (2006).

9. van Sorge, N.M. *et al.* The classical lancefield antigen of group a Streptococcus is a virulence determinant with implications for vaccine design. *Cell host & microbe* **15**, 729-740 (2014).

10. Kabanova, A. *et al.* Evaluation of a Group A Streptococcus synthetic oligosaccharide as vaccine candidate. *Vaccine* **29**, 104-114 (2010).

11. McGregor, R. *et al.* Naturally acquired functional antibody responses to group A Streptococcus differ between major strain types. *mSphere* **8**, e0017923 (2023).

12. Gallotta, M. *et al.* SpyAD, a moonlighting protein of group A Streptococcus contributing to bacterial division and host cell adhesion. *Infect Immun* **82**, 2890-2901 (2014).

13. Courtney, H.S. *et al.* Trivalent M-related protein as a component of next generation group A streptococcal vaccines. *Clin Exp Vaccine Res* **6**, 45-49 (2017).

14. Dale, J.B. *et al.* Protective immunogenicity of group A streptococcal M-related proteins. *Clin. Vaccine Immunol.* **22**, 344-350 (2015).

15. Frost, H.R. *et al.* Analysis of Global Collection of Group A Streptococcus Genomes Reveals that the Majority Encode a Trio of M and M-Like Proteins. *mSphere* **5** (2020).

16. Frost, H.R. *et al.* Promiscuous evolution of Group A Streptococcal M and M-like proteins. *Microbiology* **169** (2023).

17. Loh, J.M.S., Lorenz, N., Tsai, C.J., Khemlani, A.H.J. & Proft, T. Mucosal vaccination with pili from Group A Streptococcus expressed on Lactococcus lactis generates protective immune responses. *Sci Rep* **7**, 7174 (2017).

18. Steemson, J.D. *et al.* Survey of the bp/tee genes from clinical group A streptococcus isolates in New Zealand - implications for vaccine development. *J Med Microbiol* **63**, 1670-1678 (2014).

19. Henningham, A. *et al.* Conserved anchorless surface proteins as group A streptococcal vaccine candidates. *J Mol Med (Berl)* **90**, 1197-1207 (2012).

20. Henningham, A. *et al.* Structure-informed design of an enzymatically inactive vaccine component for group A Streptococcus. *mBio* **4** (2013).

21. McNeil, S.A. *et al.* A double-blind, randomized phase II trial of the safety and immunogenicity of 26-valent group A streptococcus vaccine in healthy adults. *International Congress Series* **1289**, 303-306 (2006).

22. McNeil, S.A. *et al.* Safety and immunogenicity of 26-valent group A streptococcus vaccine in healthy adult volunteers. *Clinical Infectious Diseases* **41**, 1114-1122 (2005).

23. Pastural, E. *et al.* Safety and immunogenicity of a 30-valent M protein-based group a streptococcal vaccine in healthy adult volunteers: A randomized, controlled phase I study. *Vaccine* **38**, 1384-1392 (2020).

24. Frost, H.R. *et al.* Immune Cross-Opsonization Within emm Clusters Following Group A Streptococcus Skin Infection: Broadening the Scope of Type-Specific Immunity. *Clin Infect Dis* **65**, 1523-1531 (2017).

25. Nordstrom, T. *et al.* Enhancing Vaccine Efficacy by Engineering a Complex Synthetic Peptide To Become a Super Immunogen. *J Immunol* **199**, 2794-2802 (2017).

26. Pandey, M. *et al.* Physicochemical characterisation, immunogenicity and protective efficacy of a lead streptococcal vaccine: progress towards Phase I trial. *Sci Rep* **7**, 13786 (2017).

27. Brandt, E.R. *et al.* Opsonic human antibodies from an endemic population specific for a conserved epitope on the M protein of group A streptococci. *Immunology* **89**, 331-337 (1996).

28. Sekuloski, S. *et al.* Evaluation of safety and immunogenicity of a group A streptococcus vaccine candidate (MJ8VAX) in a randomized clinical trial. *PloS one* **13**, e0198658 (2018).

29. Pandey, M.A.W.M.A.H.J.A.G.M.A.B.M. Long-term antibody memory induced by synthetic peptide vaccination is protective against Streptococcus pyogenes infection and is independent of memory T cell help. *Journal of immunology (Baltimore, Md. : 1950)* **190**, 2692--2701 (2013).

30. Whitcombe, A.L. *et al.* An eight-plex immunoassay for Group A streptococcus serology and vaccine development. *J Immunol Methods* **500**, 113194 (2022).

31. Hanson-Manful, P. *et al.* The novel Group A Streptococcus antigen SpnA combined with bead-based immunoassay technology improves streptococcal serology for the diagnosis of acute rheumatic fever. *J Infect* **76**, 361-368 (2018).
